# Experimental evaluation of AI-driven protein design risks using safe biological proxies

**DOI:** 10.1101/2025.05.15.654077

**Authors:** Svetlana P. Ikonomova, Bruce J. Wittmann, Fernanda Piorino, David J. Ross, Samuel W. Schaffter, Olga Vasilyeva, Eric Horvitz, James Diggans, Elizabeth A. Strychalski, Sheng Lin-Gibson, Geoffrey J. Taghon

**Affiliations:** National Institute of Standards and Technology; Gaithersburg, 20899, USA; Microsoft, Office of the Chief Scientific Officer; Redmond, 98052, USA; Johns Hopkins University; Baltimore, 21218, USA; International Gene Synthesis Consortium, South San Francisco, 94080, USA; Twist Bioscience; South San Francisco, 94080, USA

## Abstract

Advances in machine learning are providing leaps forward for beneficial applications of protein engineering, while also raising concerns about biosecurity. Recently, Wittmann *et al.* described an *in silico* pipeline of generative AI tools to reformulate sequences of concern (SOCs) as synthetic homologs that may evade detection by biosecurity screening software (BSS) used by nucleic acid synthesis providers. Experimental testing of synthetic homologs is required to ascertain the true severity of this vulnerability. We present a generalizable framework to assess biosecurity risk consisting of testing, evaluation, validation, and verification (TEVV) of AI-assisted protein design (AIPD). We determine that common AIPD models in use at the time this study was initiated (early 2024) are not yet powerful enough to reliably rewrite the sequence of a given protein, while both maintaining activity and evading detection by BSS.

---

The capabilities of AI-assisted protein design (AIPD) increase the speed and expand the scope of protein engineering. Recent examples include the generation of high-affinity SARS-CoV-2 antibodies 200 times faster than traditional modeling methods and the creation of several serine hydrolases with structures unobserved in nature (*1*, *2*). AIPD tools have also improved our ability to design and discover protein sequences that perform a desired function on par with or exceeding natural proteins (*3*, *4*). While the continuing development of this technology is important for beneficial applications, concerns have been raised about misuse.

A particular biosecurity concern is the potential for such actors to use AIPD tools to rewrite protein sequences to retain activity but evade sequence-centric biosecurity screening software (BSS) used by DNA synthesis companies. BSS software is intended to identify potential sequences of concern (SOCs) in customer orders, triggering follow-up to prevent acquisition of hazardous proteins without a legitimate use case. Current BSS workflows, almost all of which depend on similarity searches against databases of natural sequences to identify SOCs, will become less robust in a future of increasingly sequence-diverse AI-generated orders (*5*). Indeed, Lauko *et al*.’s serine hydrolase work shows that functional proteins designed using AI may look nothing like those found in nature, thus reducing the strength of current best match screening approaches (*2*, *6*). A red teaming study by Wittmann *et al*. demonstrated that the BSS used by many companies *circa* 2023 failed to detect a significant portion of SOCs deliberately obfuscated via an AIPD pipeline (*7*).

Despite mitigation of the vulnerability reported in Wittmann *et al*., concerns about its severity remain due to the lack of experimental validation for AI-generated protein designs. This highlights a key question of whether AI can accurately predict protein activity from sequence or structure. Powerful protein structure prediction models, such as AlphaFold and RoseTTAFold, excel precisely because they are trained using experimental structural data—the very same type of information they aim to predict (*8*, *9*). In contrast, protein activity prediction faces a fundamental data gap: AIPD models must extrapolate functional properties from sequence and structural training data encoding complex biochemical interactions. For example, many AIPD workflows rely on *in silico* metrics to determine likelihood of activity. Synthetic protein homologs with folds more similar to the template are assumed to be more likely to retain activity of the template. To begin to address this gap between predicted and experimental data, we employed Wittmann *et al*.’s AIPD pipeline and developed an experimental validation framework, enabling us to investigate how *in silico* metrics correspond with empirical activity measurements of synthetic protein homologs.

## Study considerations and process

To perform experimental validation safely and responsibly, we chose safe proxy protein targets rather than SOCs. Our testing, evaluation, validation, and verification (TEVV) framework used proteins of different length and classes of functional complexity to validate AI prediction through measurements of protein activity. These measurements and our validation framework can inform both future AIPD models and biosecurity strategy, and allowed us to begin to answer the outstanding question: **Is it feasible to reliably rewrite the sequence of a known protein beyond identification by BSS using open-source AIPD models, while preserving protein activity in an experimental context?**

### Target proposal and rationale

To define a broad benchmark for rewriting protein sequences using AIPD, we selected three initial protein targets assigned to basic, moderate, or advanced “difficulty classes”. We consider a protein active if it expresses, folds correctly, and interacts appropriately. We assume the difficulty of preserving protein activity scales with increasing sequence length and functional repertoire. Because the AIPD models we used generate sequences one amino acid at a time, the chance that a generated amino acid could result in an inactive final protein increases with sequence length. Similarly, as a protein’s functional repertoire expands and its complexity grows, the biophysical constraints that govern its behavior become increasingly stringent, making the generation process more susceptible to errors. For example, SH3 and PDZ domains, which facilitate binding, are known for their general tolerance to substitutions in ligand recognition regions (*10*, *11*). In contrast, the activity of metabolic decarboxylase enzymes, which coordinate multiple interaction steps, is more acutely sensitive to amino acid changes (*12*). We therefore anticipated that preserving a protein’s active state after rewriting the sequence using AI becomes increasingly challenging for longer or more functionally diverse protein sequences.

To select basic, moderate, and advanced protein targets, we considered candidates with known key residues and published activity assays. For the basic class, we selected the third PDZ (PDZ3) domain from the human postsynaptic density protein 95 (PSD95/DLG4, UniProt: P78352). Binding between PDZ3 and cysteine-rich PDZ-binding protein (CRIPT, UniProt: Q9P021), one of its protein ligands, can be approximated as a classical lock-and-key interaction (*13*). In the moderate class, we selected orotidine 5’-phosphate decarboxylase from *Saccharomyces cerevisiae* (baker’s yeast), known more commonly by its encoding gene URA3 (UniProt: P03962). As an enzyme, URA3 adds the challenge of mechanically or chemically lowering the activation energy of a chemical reaction (*14*). To represent the advanced class, we selected the T7 bacteriophage RNA polymerase (RNAP) (UniProt: P00573). T7 RNAP is an enzyme often used for *in vitro* transcription in cell-free expression (CFE) systems, due to its monomeric nature, efficiency, and fast rate of messenger RNA (mRNA) production (*15*). Active RNAP synthetic homologs must not only fold with the proper geometry and surface charge distribution to localize to a specific DNA promoter sequence, but also coordinate binding of multiple substrates and motion along DNA. These three protein targets offer a suitable scale of increasing sequence length (84, 267, and 883 amino acids, respectively) and complexity with which to experimentally benchmark AIPD capability.

### Synthetic homolog generation

For each protein, we followed the process used by Wittmann *et al*. to generate synthetic homologs (*7*) and assess their likelihood of activity based on *in silico* metrics. Briefly, we first acquired a single reviewed sequence from the UniProt database that we termed wildtype. We next identified residue positions key to each protein’s function using information gathered from structural analyses, literature, UniProt, or the Braunschweig Enzyme Database (BRENDA) (table S1) (*16*, *17*). For proteins without N-terminal fusion to other protein fragments for activity assays (URA3 and T7 RNAP), we required that sequences started with methionine to initiate protein translation. We subsequently generated synthetic homologs using three protein sequence generative models (ProteinMPNN, EvoDiff-MSA, and EvoDiff-Seq) (*18*, *19*) with four different constraints: hold the key positions constant (table S1), as well as those within 500 pm (5.0 Å) in a representative crystal structure; hold the key positions constant, as well as those within 250 pm (2.5 Å); hold only the key positions constant; and, no constraints beyond sequence length. We generated 1000 sequences for each combination of model and constraint, resulting in 12000 synthetic homologs per protein target. For each synthetic homolog, 200 predicted structures were then used to calculate the *in silico* metrics ΔpLDDT (change in predicted local distance difference test) and TM-score (template modeling score) versus wildtype. Because our goal was to determine the validity of *in silico* metrics for evaluating the likelihood of activity of potential SOCs, we sampled the entire pool of generated synthetic homologs, rather than selecting only the highest-scoring sequences. This places the study in a different category than a typical engineering campaign; we sample from the entire distribution rather than filtering to highest scores. Our wide sampling also enabled us to test activities of synthetic homologs with different levels of sequence obstruction. For each protein target, we grouped synthetic homologs into 10 clusters based on the *in silico* metrics, then randomly selected 10 synthetic homologs per cluster, creating a representative set of 100 sequences per target protein (Fig. 1). We evaluated each of these representative sets (300 proteins total) for activity.

**Fig. 1.**
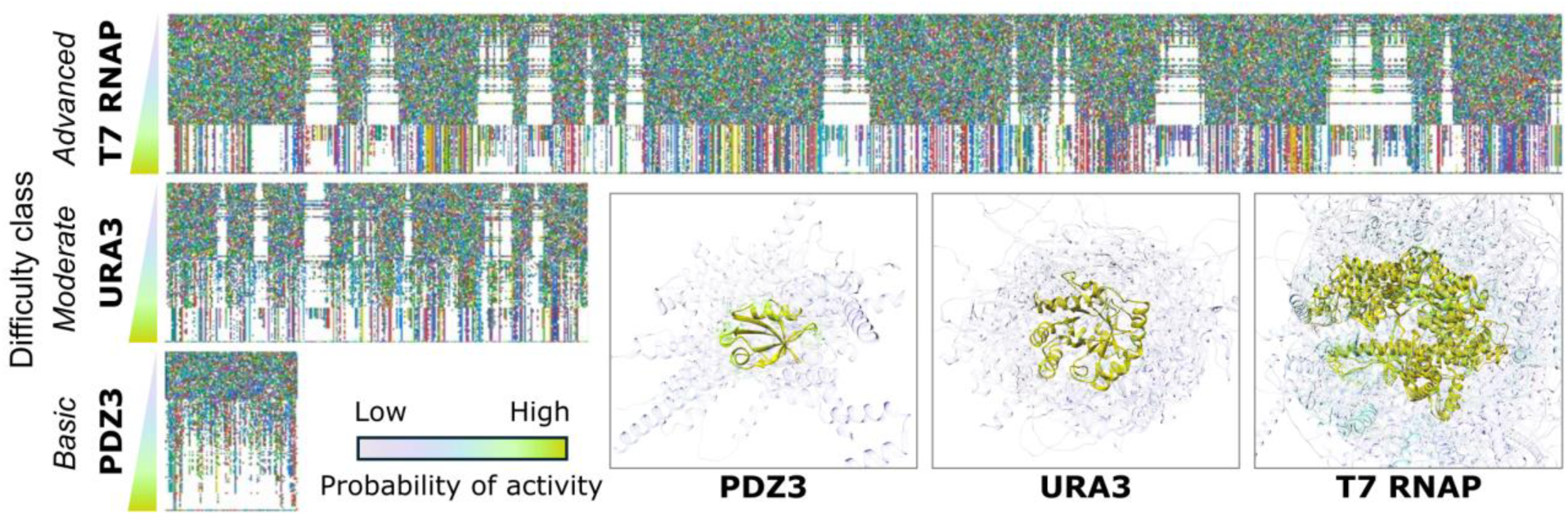
Generation of synthetic homologs of wildtype protein targets. To collect measurements to compare against AIPD predictions, we created 300 synthetic homologs spanning three difficulty classes: basic (PDZ3, n = 100), moderate (URA3, n = 100), and advanced (T7 RNAP, n = 100). Sequence-alignment plots for each protein (left) show the number and diversity of sequence substitutions increasing as AI-predicted probability of activity decreases. Each sequence is a row of pixels, where each color represents a different amino acid (*35*). Wildtype sequences are displayed on the bottom row, and a colored pixel above the wildtype sequence signifies an amino acid change from wildtype sequence. White regions of the plots represent areas of sequence conservation. For each protein target, aligned ribbon diagrams of the predicted structures for all sequences (bottom right) illustrate the increasing structural complexity of the protein targets. Structures were colored according to their AI-predicted probability of activity: lowest (transparent purple) to medium (semitransparent cyan) to highest (opaque yellow). Synthetic homologs with the highest probability of retaining activity appeared as a yellow cluster very similar to their respective wildtype structures.

### Creating a TEVV framework

To quantify the predictive capabilities of AIPD in the laboratory, we created a replicable TEVV framework. First, we developed effective activity assays for each target in appropriate model systems, drawing on existing methods and our experience with bacteria, yeast, and CFE systems (*20*–*22*). After we optimized these assays using control variants, we proceeded to test the synthetic homologs. These primary assays were intended to be rapid, informative, and self-contained, making feasible the processing of 100 sequence-diverse synthetic homologs per protein target (Fig. 2). Importantly, design of the primary assays also avoided the burdensome task of developing multiple purification protocols per protein target due to the sequence dissimilarity between synthetic homologs. Synthetic homologs were labeled as active or inactive based on a statistical threshold for each protein, and the labels were compared to the AIPD predictions. When possible, we validated the observed synthetic homolog activity using orthogonal measurements. Finally, we performed rudimentary BSS on the sequences by searching for the best match in NCBI sequence databases to determine whether the identity of the protein target remained detectable after sequence rewriting (*23*).

**Fig. 2.**
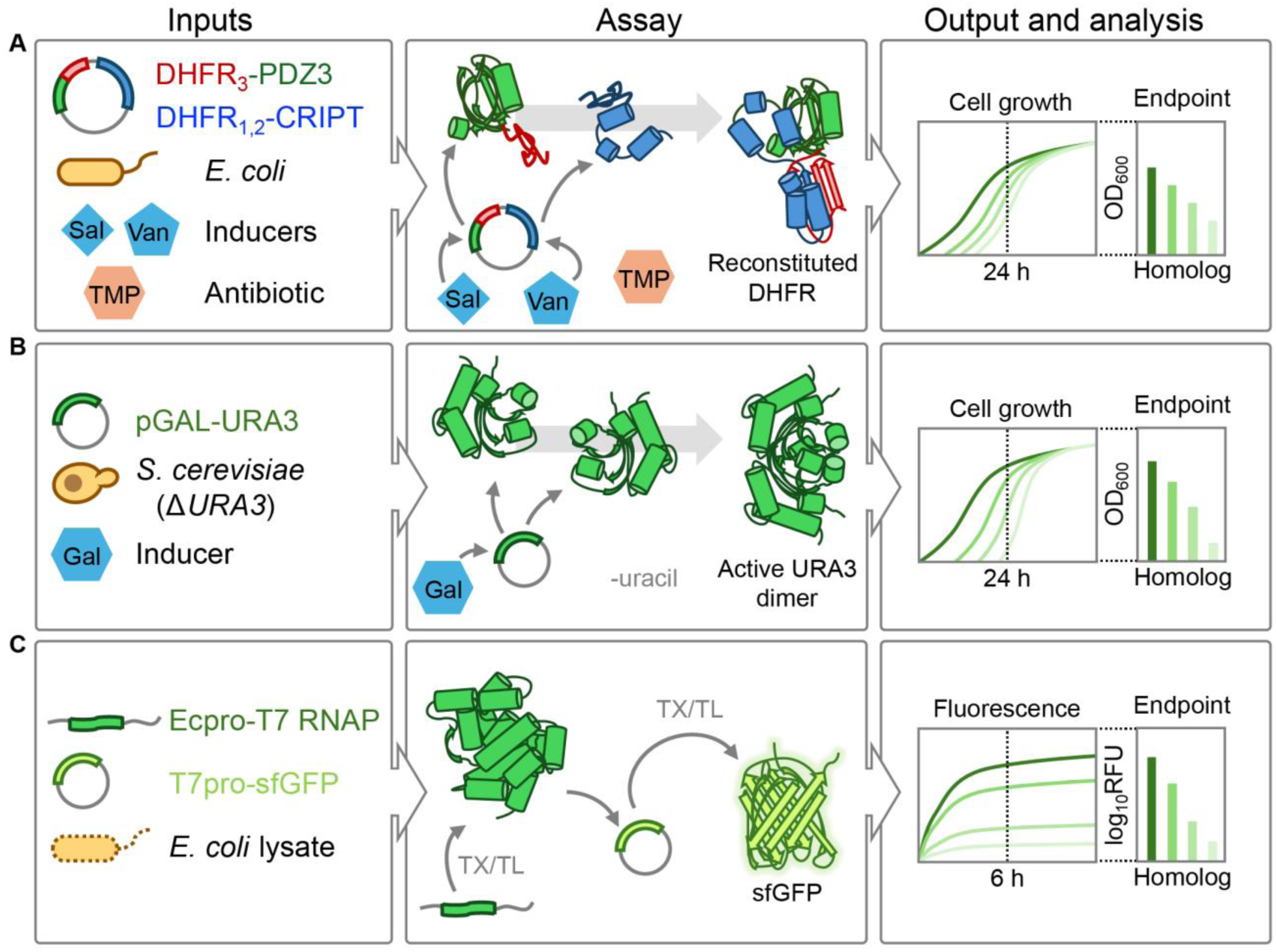
Primary assays for activity measurement of PDZ3, URA3, and T7 RNAP synthetic homologs. (**A**) PDZ3-CRIPT binding caused reconstitution of a split dihydrofolate reductase (DHFR) enzyme in *E. coli*, conferring resistance to trimethoprim (TMP). Cell growth in media with TMP was measured as an indicator of PDZ3-CRIPT binding. The assay components were expressed using individual inducer compounds (salicylate/Sal, vanillic acid/Van), allowing for tuning of the system. (**B**) The URA3 enzyme catalyzes formation of the essential metabolite uridine monophosphate. In the absence of uracil, the chosen yeast host strain grew only with an active URA3 enzyme. The enzyme was expressed from a plasmid with a galactose-inducible promoter, and cell growth in uracil-free media was measured as a reporter of URA3 activity. (**C**) T7 RNA polymerase (T7 RNAP) is commonly used in cell-free expression (CFE) systems; in this system, T7 RNAP served as both an expression target and a functional component. Linear DNA encoding T7 RNAP was transcribed and translated into protein, and T7 RNAP transcribed superfolder GFP (sfGFP) mRNA from a target plasmid. The sfGFP was then translated by the CFE system, resulting in green fluorescence as a measurable reporter of T7 RNAP activity. All assay measurements were normalized to the response of the associated wildtype proteins and negative controls.

## Results

### PDZ3

PDZ domains, such as PDZ3, are ubiquitous natural protein–protein interaction domains that bind to protein C-terminal peptides. To avoid the time- and resource-consuming task of purifying 100 synthetic homologs of PDZ3 required to directly measure binding, we initially assessed activity using a high-throughput primary cellular assay in *Escherichia coli*. We employed a split-dihydrofolate reductase (DHFR) protein complementation assay to measure PDZ3-CRIPT interactions (*24*, *25*). PDZ3 and its natural ligand CRIPT were each fused to a DHFR fragment. When PDZ3 and CRIPT interact, the DHFR fragments come into proximity and reconstitute an active enzyme. Active DHFR confers resistance to the antibiotic trimethoprim (TMP) and allowed cell growth in the presence of TMP. Thus, we measured cell growth to quantify binding activity (Fig. 2A). We first tested the cellular assay using four PDZ3 control variants: wildtype, two PDZ3 variants reported to have reduced affinity to CRIPT (A376R and Y392I), and a non-binding variant with the CRIPT sequence omitted (*11*). Results demonstrated clear differences among control variants: reduced-affinity controls showed reduced activity compared to wildtype, and the non-binding control exhibited very low activity (fig. S1). Upon testing the synthetic homologs, we found 29 of 100 were active above our threshold (*see Materials and Methods*), with 14 displaying activity equal to or exceeding wildtype PDZ3 (Fig. 3, Fig. 4, and fig. S2). Surprisingly, Homologs 70 and 77, with minimal structural and sequence similarity to wildtype, exhibited low but above-threshold responses. These sequences were generated using only the length of PDZ3 as a constraint, and active synthetic homologs were unexpected for such an otherwise unconstrained generation.

**Fig. 3.**
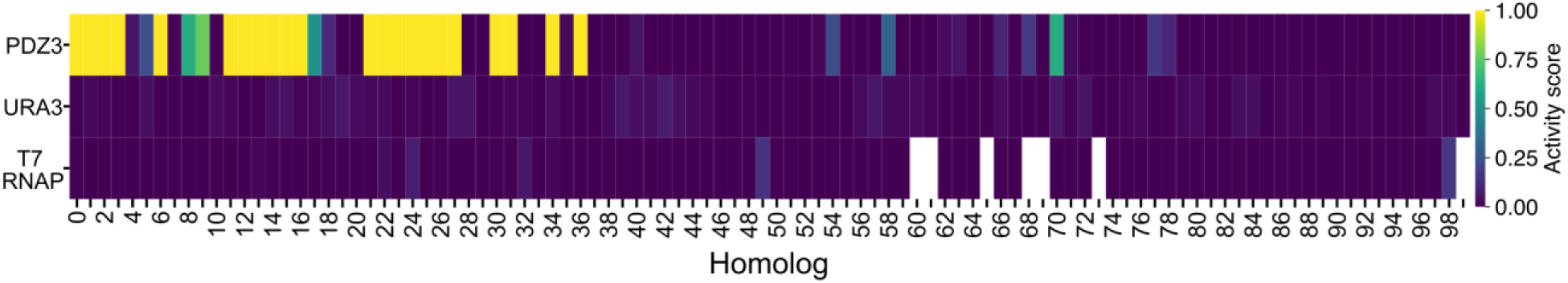
Activity scores for each protein target show no activity for the majority of synthetic homologs. For each protein target, primary assay activity scores were determined by normalizing the activity of a synthetic homolog to the relevant wildtype and negative controls. Only a subset of PDZ3 synthetic homologs demonstrated substantial activity. Synthetic homologs are numbered by their likelihood of activity according to AIPD metrics. Seven T7 RNAP synthetic homologs (white) could not be synthesized or cloned.

**Fig. 4.**
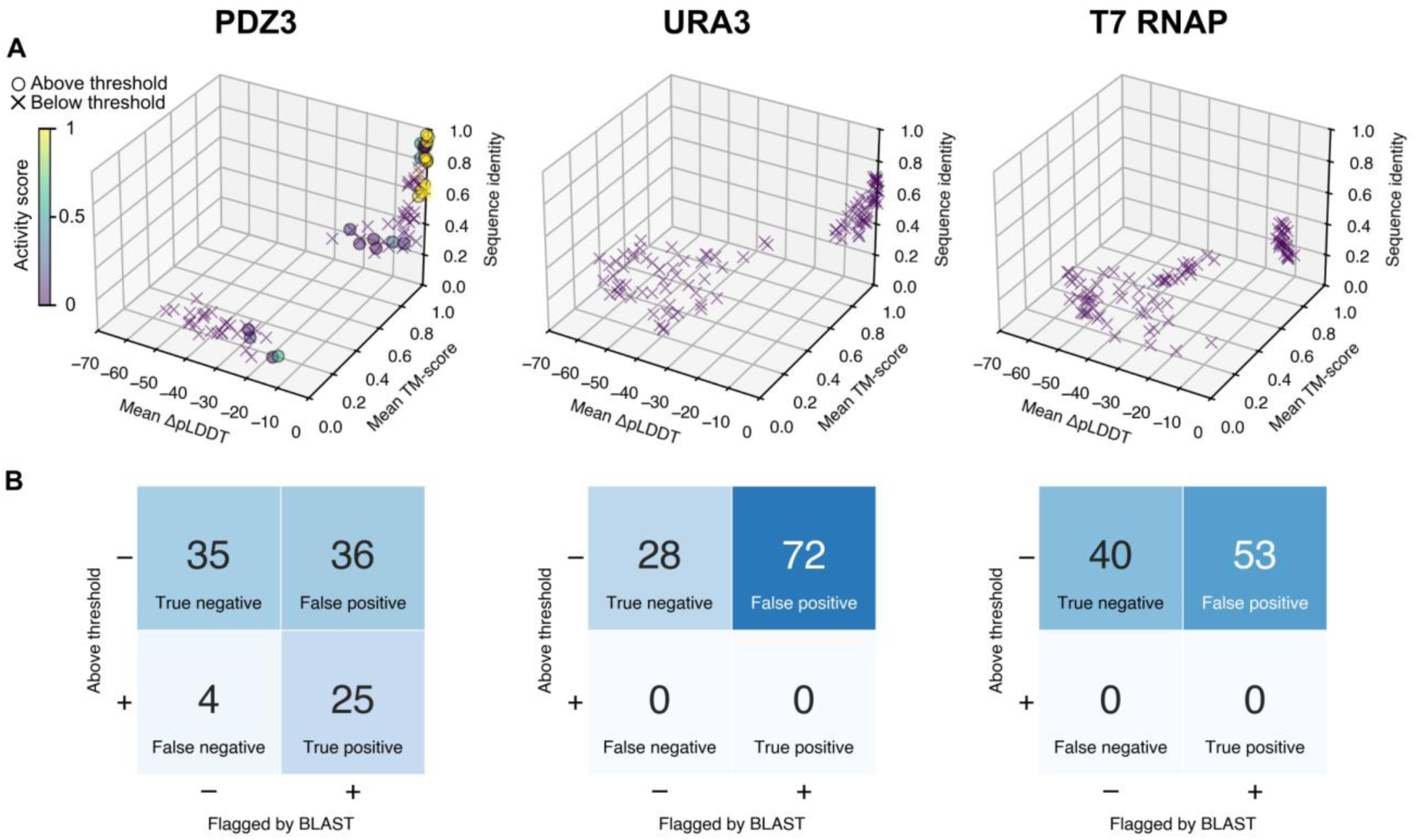
Comparison of AIPD-calculated metrics to the measured activity of synthetic homologs and potential for sequence screening evasion. (**A**) Plots show synthetic homologs of each protein as points in a coordinate space defined by the mean ΔpLDDT, mean TM-score, and sequence identity compared to wildtype sequences (PDZ3: n = 100, URA3: n = 100, T7 RNAP: n = 93). Points are colored by activity score calculated as the activity normalized to wildtype and negative controls. Active synthetic homologs are plotted as circles, inactive synthetic homologs are plotted as crosses. The upper right corner of each plot contains synthetic homologs predicted by the model as the most likely to retain activity (highest structural prediction confidence, fold similarity to wildtype, and sequence identity), while the lower left corner contains the synthetic homologs predicted as the least likely to retain activity. (**B**) To determine if active synthetic homologs were effectively sequence obfuscated and therefore able to evade sequence screening, we submitted them to NCBI BLAST and evaluated the best match entry. Analogous to how BSS identifies obfuscated SOCs, if the best match contained a keyword referring to the wildtype sequence, the synthetic homolog was designated as “flagged”—indicating it was identifiable as derived from wildtype. Thus, synthetic homologs that were experimentally active and flagged by BLAST were “True positive.”

To investigate the specificity of the observed activity of the synthetic homologs, we modified the cellular assay to reevaluate selected synthetic homologs with either activity above wildtype or activity above the threshold despite low sequence similarity to wildtype. We replaced the CRIPT C-terminal peptide, which binds to PDZ3, with control peptides previously reported to exhibit low affinity to PDZ3 in *in vitro* assays (*see Materials and Methods*) (*26*). For these control peptides, we chose one natural peptide, Nat18, and one synthetically derived peptide, Syn15. To confirm that the C-terminal modifications would not disrupt the backbone, we performed structural modeling and alignment of these modified CRIPT constructs to wildtype CRIPT (fig. S3). The modified cellular assay confirmed no activity for wildtype PDZ3 with Nat18, as anticipated, but displayed some activity for wildtype PDZ3 with Syn15 (fig. S4). While most of the tested synthetic homologs, including Homologs 70 and 77, similarly exhibited no activity with Nat18 and some activity with Syn15, several synthetic homologs were active with both peptides. These differing activities among the synthetic homologs indicate a possible change in the specificity of PDZ3 binding (fig. S4). We conclude that activity observed in both the primary and modified cellular assays for some of the synthetic homologs may be attributed to nonspecific interactions separate from binding between the CRIPT C-terminus and PDZ3.

To further probe the PDZ3 binding specificity, we tested a subset of the synthetic homologs with an *in vitro* binding assay using biolayer interferometry (BLI) (*see Materials and Methods*). We initially sought to purify synthetic homologs with and without the DHFR fragment from the cellular assays. However, we were unable to achieve the yield or purity needed for BLI measurements with DHFR fragment fusions. We successfully purified wildtype PDZ3 and Homologs 9, 11, and 36, without DHFR fragments (fig. S5). We then evaluated binding activity with ligand peptides immobilized on the biosensor tip (*see Materials and Methods*) (*27*). None of the purified proteins had activity with Syn15 peptide. Wildtype PDZ3, Homolog 9, and Homolog 11 showed activity with CRIPT peptide, and only Homolog 11 had activity with Nat18 peptide. Homolog 36, however, had no activity with any peptide, supporting the conclusion that some of the observed activity in the cellular assays is due to nonspecific interactions, especially for synthetic homologs with low sequence identity compared to wildtype (fig. S6). Steric hindrance may have also contributed to discrepancies between cellular assays and BLI measurements. The ligand peptides were immobilized in the BLI assay but were at the C-termini of cytosolic proteins in our cellular assays. These differences may have impacted the accessibility of the binding region and therefore the observed binding activity. Together, these results revealed that the evaluation of sequence diversity of synthetic homologs can require multiple assays.

From a sequence screening perspective, ∼86 % of the synthetic homologs labeled experimentally active returned a best match via NCBI BLAST to a known PDZ domain-containing protein (True positive in Fig. 4B). However, protein annotation detail affected our simple approach to sequence identification. For example, BLAST matched multiple inactive synthetic homologs to the PDZ-containing Disks large (DLG) protein family, but the “PDZ” string did not appear in the GenBank entries (table S2). Therefore, these synthetic homologs were incorrectly classified as true negatives. Another notable result was the apparent activity of Homolog 70. This synthetic homolog has low sequence similarity to known PDZ domains, as evidenced by a lack of BLAST results, and its predicted structure bears no resemblance to PDZ folds (table S2 and fig. S7). We conclude our observed activity of Homolog 70 arose from interactions other than those between PDZ3 and CRIPT. This further highlights the importance of multiple orthogonal methods to validate specific protein interactions. Importantly, none of our orthogonally-validated, active synthetic homologs would have evaded minimal recommendations for BSS (*6*).

### URA3

URA3 is a yeast auxotrophic enzyme commonly used as a selection marker. Yeast cells grow in media without uracil only when expressing active URA3 (*28*). Therefore, cell growth can provide an indirect measure of URA3 enzyme activity (Fig. 2B) (*29*). As with growth-based measurements for PDZ3 activity, this cellular assay avoided onerous purification of synthetic homologs. The Δ*URA3* yeast strain was transformed with a galactose-inducible plasmid expressing URA3 controls or synthetic homologs (*28*, *30*). Expression of URA3 in uracil-free media with galactose restored cell growth. We first tested control variants: wildtype and two URA3 variants with reduced activity (R235A and Q215A) (*12*). We confirmed the reduced activity of R235A and undetectable activity of Q215A, as compared to wildtype URA3. Our tests of the 100 synthetic homologs yielded no measurable activity, likely due to sequence rewriting on the dimer interface and catalytic site (Fig. 3, Fig. 4, and fig. S8). Q215A exemplified how a single amino acid substitution in the catalytic site can effectively abolish activity—among our synthetic homologs, the closest sequence to wildtype URA3 (Homolog 2) contained 82 substitutions.

Despite this sequence distance, the identification of URA3 synthetic homologs with BLAST screening was remarkably effective (Fig. 4B). 72 of 100 sequences returned a form of “decarboxylase”. Curiously, Homolog 76 was the least distant sequence to return no BLAST match, while more distant Homologs 89 to 94 were matched to an orotidine 5-phosphate decarboxylase enzyme, either from a yeast or a bacterium (table S2). Sequence identity to wildtype URA3 for Homologs 89 to 94 ranged from ∼(12.7 to 19.9) %. This finding suggests key residues used as explicit (user-supplied) and implicit (model-learned) constraints provide a strong signature identifiable by BSS, despite the extreme rewriting that resulted in loss of activity.

### T7 RNAP

Given the widespread use of T7 RNAP in CFE systems, we began by testing synthetic homologs for activity in this context. To mitigate the impact of T7 RNAP cytotoxicity during DNA synthesis and purification, controls and synthetic homologs were cloned into a plasmid with a mutated, nonfunctional promoter sequence. Of the 100 synthetic homologs, 93 could be synthesized. For the primary CFE assay, we prepared linear templates from these plasmids using high-fidelity PCR with a 5’ primer that repaired the promoter sequence. These templates were then added to a CFE system optimized for linear DNA (*31*), which also contained a plasmid encoding a superfolder GFP (sfGFP) reporter gene under control of a T7-specific promoter. The linear templates were transcribed and translated into T7 RNAP, and active polymerase then transcribed the sfGFP gene to produce a fluorescence signal to measure polymerase activity (Fig. 2C and fig. S9). We identified no statistically-significant activity for synthetic homologs (*see Materials and Methods*, Fig. 4, and fig. S10). However, we observed notable signal of ∼10 % of wildtype for Homolog 98 (sequence identity ≌ 6.5 %, mean TM-score = 0.22) (fig. S10A). This synthetic homolog resulted from an effectively random sequence generation with only sequence length as a constraint, so the unusual result prompted further testing.

Variability in the replicates of the primary CFE assay suggested introduction of contaminants or sequence errors from the PCR process or degradation of the linear DNA templates. Thus, we selected a subset of synthetic homologs and proceeded with a plasmid template variation of the CFE assay. Nine synthetic homologs were successfully cloned, including Homolog 98; none displayed improved signal for a plasmid compared to linear template (fig. S10C). To control for sequence-dependent transcript stability and transcription rates, we performed CFE assays with purified mRNA produced by *in vitro* transcription. We quantified and normalized the mRNA concentrations, then added them to the assay, bypassing transcription in the CFE system. We observed no improvement in our measurement as compared to DNA templates, and only Homolog 98 yielded a measurable fluorescence output (fig. S10D). Finally, we assayed a subset of synthetic homologs, including Homolog 98, in an *E. coli* cellular assay using inducible cassettes and a fluorescent reporter; we observed no synthetic homolog activity (fig. S11A). Therefore, we hypothesize that the spurious activities observed in the primary CFE assay arose as measurement artifacts.

BSS testing provided more conclusive results. We found that a significant proportion of T7 RNAP synthetic homologs were unidentifiable by BLAST (37 of 93 assayed sequences, ∼40 %), perhaps due to the length of the sequence and lower overall synthetic homolog sequence identity. However, 53 of 56 identified sequences (∼95 %) still matched to “RNA polymerase” (Fig. 4B). Therefore, even for long and highly dissimilar protein sequences in the advanced difficulty class, best match and other sequence-based BSS strategies remain useful to determine the intent behind a sequence order. Although these T7 RNAP synthetic homologs did not demonstrate activity in assays, BLAST was often able to detect their native protein source.

## Discussion and recommendations

### AIPD TEVV is difficult

Despite encouraging steps towards the experimental validation of AIPD, multiple formidable obstacles remain. Scientists with the computational, biological, and technical expertise required to create and execute validation assays are difficult to assemble. In a machine learning research climate that discourages negative results and rewards *in silico* predictive performance, model developers are not currently incentivized to submit AIPD generations for laboratory TEVV (*32*). Groups with access to laboratory resources and the desire to perform TEVV studies are likewise disincentivized to characterize sequences generated with a previously published AIPD pipeline, as this type of work may not be seen as novel or innovative, despite its obvious importance. Even with the appropriate infrastructure and team in place, experimental validation can be prohibitively expensive and time-intensive, which may discourage researchers and funders. The development, validation, and execution of activity assays often requires a substantial period of time, typically ranging from months to years. In contrast, training new generations of AIPD models can be completed much more rapidly. This significant disparity in timelines may lead researchers to prioritize timely publication over experimental validation (*33*).

Regarding biosecurity, working with SOCs requires special oversight, safety practices, and logistics, adding additional layers of risk and experimenter aversion. These challenges discourage the evaluation of AIPD models in the laboratory and prevent determining the actual risks of misuse. Our solution presented here is collaborative, low-risk TEVV using safe proteins as proxies for SOCs. Delegation of these tasks across different organizations reduced the time needed to complete a TEVV study, and we eliminated risk associated with SOCs by choosing biologically interesting, benign proteins for this study. We distributed the workload across partners: an industry-based team provided modeling and nucleic acid synthesis resources, and a government laboratory provided high-throughput measurement infrastructure, assay design, and experimental execution. Then, all parties assessed the data and drew conclusions to provide recommendations to the research and security communities.

### Implications for biosecurity

Experimental evaluation of AI-driven protein design was not a straightforward process, from protein target proposal to DNA synthesis, assay optimization, and result validation. To increase our chances of success, we chose proteins and assay systems with which we already had some experience. This may not represent the work of cutting-edge researchers or malicious actors with limited resources working with less-studied toxic proteins; such efforts would prove more challenging. Our assays and control sequences were already published and required only protocol optimization and adaptation to liquid handling robotics. If we instead had to create, validate, and optimize new assay methods and controls, the time to completion of this study may have increased significantly. Even so, experimental validation and data collection were indeed the largest part of our timeline, requiring ∼75 % of the total duration.

Despite extensive planning, experimentation also added unforeseen constraints following the start of the project. We intended to assay additional proteins, such as the *Mycobacterium tuberculosis* toxin-antitoxin system. However, the control sequence variants proved to be too cytotoxic for DNA synthesis and cloning, even with a minimal leakage expression system (*34*). While we likely could have mitigated issues with these constructs through investing more time and effort, additional protein targets were excluded from the final project. Even after receiving sequence-verified DNA, all assays required multiple rounds of adjustments and tuning. Any attempt at AIPD TEVV needs to repeat this careful process, posing a further technological barrier.

In addition, we pursued orthogonal validation of active synthetic homologs to determine off-target effects or metabolic interference. Even in the basic class of PDZ3, orthogonal measurements sometimes did not match results from the cellular assays. If the benefit provided by AIPD is such that a potential bad actor might have to spend months in a well-equipped laboratory to verify the activity of basic class sequences that were identifiable by rudimentary BSS, we conclude that the risk of AIPD misuse is currently minimal. Furthermore, the cost of producing sequence perfect synthetic homologs, although decreasing, still limited us to analyzing 100 synthetic homologs per protein target, despite our experimental throughput capability of 10^5^ protein sequences (*20*). Our experiences support the view that the use of AI tools for the design, construction, testing, and deployment of engineered proteins remains a very challenging and resource-intensive process, especially without large sets of training data, a skilled team, substantial funding, and significant time.

### Strengths of the AIPD pipeline

We did not observe reliable prediction of synthetic homolog activity by the AIPD pipeline for moderate or advanced difficulty protein targets. However, we are surprised by the apparent activity of synthetic homologs in the cellular assay for our basic protein target, PDZ3. To assess the ability of AIPD *in silico* metrics to predict PDZ3 activity, we conducted a receiver operating characteristic (ROC) analysis. This analysis allowed us to evaluate how effectively these metrics distinguish between active and inactive synthetic homologs: ΔpLDDT (area under curve (AUC) = 0.733), followed by mean TM-score (AUC = 0.725) and sequence identity to wildtype (AUC = 0.720) (fig. S12). The similar AUC values indicate that structural metrics provide approximately the same predictive capability as sequence metrics for activity preservation of the structurally static PDZ3 domain. Indeed, all three metrics show strong covariance (average Spearman’s *ρ* = 0.91). These relationships support the conclusion that, although sequence identity had the lowest AUC, BLAST screening easily identified a majority of synthetic homologs even at high sequence and structural distance from wildtype (Fig. 4). We also observed the predicted structure of PDZ3 is particularly tolerant of sequence substitutions, with the mean TM-score declining abruptly at 74 substitutions from wildtype (fig. S13).

Our success rate for generating active PDZ3 synthetic homologs should be evaluated conservatively, given our low threshold for identifying a sequence as active—significant activity above the non-binding control—and our limited orthogonal validation. Overall, this AIPD pipeline can provide preliminary filtering for sequence prioritization, while experimental evaluation remains essential. AIPD does not currently offer a “magic bullet” for altering protein sequences while improving or even preserving activity.

### Weaknesses of the AIPD pipeline

For moderate and advanced classes involving dimer interfaces or dynamic movement of protein domains, the extensive sequence rewriting allowed by the pipeline appears to be too extreme to preserve activity. Despite the high mean TM-score of URA3 synthetic homologs, no synthetic homolog displayed activity above the threshold. However, a majority of these sequences were still identified correctly by BLAST as matches to orotidine-5’-phosphate decarboxylase, which we refer to as URA3 in this work (Fig. 4B). In the advanced class of T7 RNAP, we observed no verifiable activity in primary assays or orthogonal measurements. Similar to URA3, the majority of synthetic T7 homologs were identified correctly by BLAST (Fig. 4B). Even with the limited scope of three proteins and 293 synthetic homologs, our investigation offers preliminary evidence that avoiding BSS detection requires synthetic homologs with sequences far from the wildtype protein, which would typically cause activity loss despite structural similarity.

Together, our results have direct implications for both protein engineers and those seeking to build systems to defend against potential misuse of AIPD. We offer a provisional answer to our main question: **Using open-source AIPD models, rewriting the sequence of a known protein beyond identification by BSS while preserving activity does not currently appear feasible.** Sequence identity is also revealed to be a barrier against hypothetical AIPD-enabled screening evasion, as evidenced by our successful BLAST-based identification of active synthetic homologs. Importantly, however, these conclusions do not apply to completely synthetic folds, such as the novel hydrolases mentioned previously (*2*). This concept is relevant to our *in silico* investigation of PDZ3 Homolog 70; it almost certainly lacks a PDZ3 domain yet appeared active in our cellular assay. Regardless, intentional generation of novel folds has thus far required models and teams with deep mechanistic understanding, and these scenarios require consideration exceeding the scope of our study.

### Outlook

As AIPD capabilities advance rapidly, several key opportunities for improvement emerge. First, the collection and incorporation of experimental data into model training would significantly increase AIPD models’ understanding of protein function. This effort requires setting realistic experimental goals and prioritizing proteins with well-understood functions, such as our safe proxies, which can provide valuable insights for BSS. Second, we suggest the basic, moderate, and advanced classes, along with their corresponding primary assays presented here, provide a preliminary benchmark to monitor AIPD performance. While our study confirms that generative models can recapitulate predicted structure, this does not necessarily correlate with preserving activity. Notably, when activity was preserved in generated sequences, their origin was typically identifiable through sequence homology-based searches, highlighting the importance of key residues and motifs. This finding tempers concerns about the robustness of BSS and the potential for sequence obfuscation as an evasion technique. Ultimately, we demonstrate that experimental validation of AIPD capabilities is not only essential but also achievable, with our framework offering as a blueprint for future studies.

## Acknowledgments

We thank Kinlin L. Chao and Zvi Kelman for facilitating the use of laboratory space and equipment at the Institute for Bioscience and Biotechnology Research for PDZ3 purification and in vitro testing. We thank Molly E. Wintenberg and Stephanie L. Guerra for discussions and feedback.

Certain tools, software, commercial equipment, instruments, or materials are identified to adequately specify the experimental procedure. Such identification implies neither recommendation or endorsement by the National Institute of Standards and Technology nor that the tools, software, materials, or equipment identified are necessarily the best available for the purpose. The views expressed in this publication are those of the authors and do not necessarily represent the views of the U.S. Department of Commerce or the National Institute of Standards and Technology.

## Supplementary Materials

Figs. S1 to S14

Tables S1 to S2

## Author contributions

Conceptualization: SPI, DRJ, EH, SL, GJT

Data curation: SPI, FP, DRJ, SWS, GJT

Formal analysis: SPI, FP, DRJ, SWS, GJT

Investigation: SPI, FP, DRJ, SWS, OV

Methodology: SPI, FP, DRJ, SWS, GJT

Project administration: SPI, EAS, GJT

Resources: OV, JD

Software: GJT

Supervision: EAS, GJT

Validation: SPI, FP, DRJ, SWS

Visualization: SPI, FP, DRJ, SWS, GJT

Writing – original draft: SPI, GJT

Writing – review & editing: SPI, BW, FP, DRJ, SWS, EH, JD, EAS, SL, GJT

## Data and materials availability

Data and analysis code is available at github.com/usnistgov/AIPD_TEVV.

## Materials and Methods

### Computational

#### Synthetic homolog generation

Synthetic homolog sequences were generated following the approached outlined in Wittmann *et al*. <u>(*7*)</u>.

#### Sequence alignment

The graphical sequence alignments in Figure 1 were prepared using the AliView software package <u>(*35*)</u>. Each input protein target FASTA file contained the 100 synthetic homolog sequences sorted by decreasing probability of activity and ends with the wildtype sequence. Sequences were aligned and colored by amino acid. Then, the wildtype sequence was set as the reference sequence. Only the full wildtype sequence and synthetic homolog substitutions from wildtype are plotted in the alignment graphics to highlight the trend in sequence diversity versus probability of activity.

#### CRIPT C-terminal variant modeling

##### AlphaFold3

CRIPT protein C-terminus(CRIPT: VCEKCEKKLGRVITPDTWKDGARNTTE-SGGRKLNENKALTSKKARFDPYGKNKFSTCRICKSSVHQPGSHYCQGCAYKKGICAMC GKKVLD**TKNYKQTSV**, C-terminus in bold) was modified to the following sequences from Kurakin, *et al*. for use as controls: VSASHDTIV (Syn15) and IAESLESLV (Nat18) <u>(*26*)</u>. CRIPT, CRIPT-Syn15, and CRIPT-Nat18 were submitted to the AlphaFold3 server for mmCIF model generation using the default settings <u>(*8*)</u>. The mmCIF models were then converted to PDB format for visualization using UCSF Chimera <u>(*36*, *37*)</u>.

##### Structural alignment

Model 0 for CRIPT-Syn15, CRIPT-Nat18 were aligned to the AlphaFold3 model for CRIPT (UniProt: Q9P021) using UCSF Chimera’s MatchMaker tool <u>(*36*,</u> *<u>37</u>*). Root mean square deviation (RMSD) was reported in picometers (pm) and Angstroms (Å) (fig. S3).

### Data analysis

#### Activity threshold

We classified synthetic homologs as active if their assay output significantly exceeded negative control output. To determine significance, different methods were used based on the type of data. For PDZ3 and URA3 cellular assays, synthetic homologs were considered active if their OD_600_ measurement significantly exceeded the negative control OD_600_ measurement, as determined by a one-tailed Welch’s t-test (*p<*0.05). For the T7 RNAP CFE assay, the mean log_10_RFU values and activity scores were estimated using a Bayesian hierarchical model that included both day-to-day and within-day variability. The results of the hierarchical models were also used to calculate the posterior probability that the activity score for each synthetic homolog is negative. Conditions were considered active if their calculated posterior probability was less than 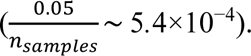 These thresholds were chosen because even minimally active sequences can be concerning—malicious actors could use natural selection and other evolutionary techniques to rapidly enhance activity.

#### Sequence screening

As a rudimentary form of BSS, we submitted each synthetic homolog sequence to NCBI BLAST to determine the best match to any existing database entries, if any. Then, the complete GenBank or other database entry was searched for a relevant keyword or phrase (“PDZ”, “decarboxylase”, “RNA polymerase”). For example, if the keyword “PDZ” was found within the best match entry for a PDZ3 sequence, we considered that sequence to be “flagged”. All words in a phrase had to be found in the entry: *i.e.*, entries containing only “RNA” or “polymerase” did not raise a flag. If there were zero BLAST matches or the keyword or phrase was not found, the sequence was not “flagged”.

#### Structure model generation in PDB format

PDB models were created using OpenFold (*38*). Models of the wildtype protein and synthetic homologs were each predicted 200 times using unique seeds.

#### AIPD metric calculation

##### Mean ΔpLDDT

The pLDDT scores for the set of wildtype and synthetic homolog models were used to calculate 200 ΔpLDDT values for each of the synthetic homolog sequences (*38*). Then, the mean was calculated:

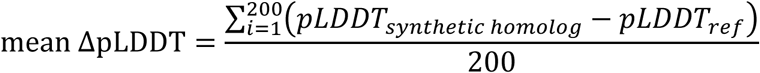

##### Mean TM-score

The template modeling (TM)-score was calculated between each synthetic homolog PDB model and each wildtype PDB model, yielding 200 TM-scores. Then, the mean was calculated:

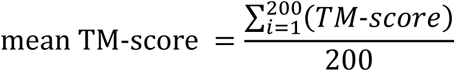

##### Percent Identity

The sequence identity between each synthetic homolog sequence and its wildtype sequence was calculated:

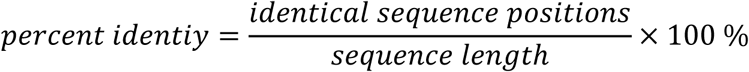

##### Probability of activity

The TM-scores, ΔpLDDTs, and sequence identities were used to define a coordinate space to map synthetic homolog sequences. A composite of the three *in silico* metrics used for clustering was calculated by: mean-centering to zero and unit-scaling each dimension of the metric space; min-max scaling between 0 and 1 for each dimension of the mean-centered, unit-scaled metric space; and, calculating the 3D Frobenius norm (Euclidean distance between origin and the point) of the point in metric space:

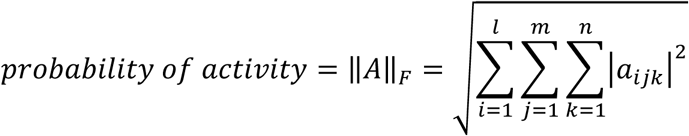

### PDB annotation with probability

Probability of activity was used to annotate the OpenFold model PDB files, overwriting “occupancy”. This allowed for simple recoloring and display of model chains in UCSF Chimera (“Depiction: Render by attribute”) (Fig. 1) (*36*, *37*).

#### PDB alignment

To produce the ribbon diagram panels in Figure 1, synthetic homolog models were aligned to each other using the UCSF Chimera software’s MatchMaker tool (*36*, *37*). Chains were then colored by probability of activity as described in “AIPD Metric Calculation” to highlight the trend in model structure versus probability.

### Assays

#### PDZ3

##### DNA constructs and plasmid transformation

Plasmids for the cellular assays were obtained from Twist Bioscience using the “pTwist Kan Medium Copy” cloning vector (table S1). The plasmid design contained bicistronic design translational control elements (BCDs) (*39*) and insulators (RiboJ and PlmJ) (*40*) to account for sequence-specific variations in expression (fig. S14). Plasmids for the *in vitro* assay were ordered from Twist Bioscience using their pET21(+) expression vector with the ribosome binding site (RBS) from pET21a(+) plasmid added. The PDZ3/DHFR_3_-PDZ3 inserts also included a 6×histidine-tag and TEV cleavage site at the N-terminus.

##### PDZ3 cellular assay

Plasmids containing DHFR_1,2_-CRIPT and DHFR_3_ fused with wildtype PDZ3, known control variants (A376R and Y392I) (*11*), or synthetic homologs were transformed into the *E. coli* Marionette-Clo strain derived from DH10*β* (*41*). For the non-binding control of wildtype PDZ3, the CRIPT sequence was removed from DHFR_1,2_-CRIPT fusion. Transformed cells were plated on LB-agar plate with kanamycin (Kan) selection. Three colonies were picked from each plate and grown overnight in M9-glycerol media (1× Difco M9 minimal salts, 0.4 % (v/v) glycerol, 0.2 % casamino acids, 100 *µ*mol/L calcium chloride, and 2 mmol/L magnesium sulfate) with 50 *µ*g/mL Kan. Cryogenic stocks were prepared by mixing overnight culture with 40 % glycerol at 1:1 volume ratio.

Before loading onto an automated liquid handling platform, the following steps were taken:

1. Overnight cultures were started from glycerol scrapings and grown in 5 mL of M9-glycerol media with 50 *µ*g/mL Kan in 14 mL snap-cap culture tubes at 37 ^◦^C, 300 rpm, for 22 h.
2. On the day of experiment, media, ligand, and antibiotic working solutions were prepared.

a. Kan was added to 400 mL of M9-glycerol media to final concentration of 50 *µ*g/mL (M9-gly-kan).
b. Antibiotic working solution (10× final concentration) with 125 *µ*g/mL trimethoprim (TMP) and 0.5 % (v/v) dimethyl sulfoxide (DMSO) was prepared using part of the M9-gly-kan media and TMP stock solution of 25 mg/mL (in DMSO).
c. DMSO was added to the rest of the M9-gly-kan media to a final concentration of 0.5 %.
d. Working ligand solutions (5× final concentration) of sodium salicylate (Sal) at 250 *µ*mol/L, and vanillic acid (Van) at 500 *µ*mol/L were prepared using M9-gly-kan with 0.5 % DMSO. (The stock solution of Sal was prepared at 100 mmol/L in water and the stock solution of Van was prepared at 100 mmol/L in ethanol).
3. The prepared media (M9-gly-kan with 0.5 % DMSO), working antibiotic solution, ligand solutions, and overnight cultures were loaded onto the automation system to begin the cellular assay.

The following processes were carried out with an automated liquid handling platform in sequential order:

1. Growth Plate 1 preparation: 450 *µ*L of M9-gly-kan media with 0.5 % DMSO was mixed with 50 *µ*L of overnight cultures (1:10 dilution of cells).
2. Growth Plate 1 measurement: Growth Plate 1 was sealed with a breathable membrane and moved to a Biotek Synergy Neo2 reader (Agilent Technology). Plates were incubated at 37 ^◦^C, with shaking, and OD_600_ measured every 5 min for 12 h.
3. Growth Plate 2 preparation: 490 *µ*L of media with and without inducers (final concentration 50 *µ*mol/L Sal and 100 *µ*mol/ Van) was mixed with 10 *µ*L of cells from Growth Plate 1.
4. Growth Plate 2 measurement: Growth Plate 2 was sealed and incubated at 37 ^◦^C, with shaking, and OD_600_ measured every 5 min for 3 h.
5. Growth Plate 3 preparation: 490 *µ*L of media with inducers and TMP (final concentration 12.5 *µ*g/mL) was mixed with 10 *µ*L of cells from Growth Plate 2.
6. Growth Plate 3 measurement: Growth Plate 3 was sealed and incubated at 37 ^◦^C, with shaking, and OD_600_ measured every 5 min for 24 h.
7. OD_600_ at end of Plate 3 measurement was used to evaluate activity.

*Metrics:* Normalized activity score was calculated as:

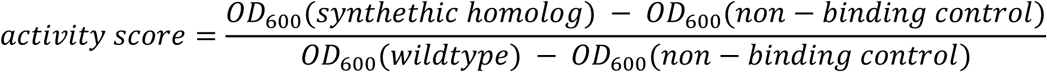

at 24 h during the Growth Plate 3 measurement. Any synthetic homologs with growth significantly exceeding the non-binding control as determined by a Welch’s one-tailed t-test (*p<*0.05) were labeled active.

#### PDZ3 in vitro assay

##### PDZ3 and synthetic homologs expression

Plasmids with PDZ3 wildtype and synthetic homologs, with and without DHFR fusions, were first transformed into BL21 Star (DE3). The following steps were taken to express proteins for purification:

1. A 10 mL starter culture was grown overnight at 37 ^◦^C, 225 rpm and used to inoculate 1 L LB broth (10 g/L sodium chloride, 5 g/L yeast extract, and 10 g/L tryptone) with 100 *µ*g/mL ampicillin in a 4 L baffled flask.
2. The 1 L cultures were grown to an OD_600_ of 0.5 to 0.8 at 37 ^◦^C, (200 to 225) rpm.
3. Cultures were cooled to room temperature, induced with 1 mmol/L of isopropyl *β*-D-thiogalactopyranoside (IPTG), then expressed overnight at 18 ^◦^C, (200 to 225) rpm.
4. Cell cultures were split into two half-volumes and harvested by centrifugation at 5000 rpm, 4 ^◦^C, for 20 min. Each pellet was frozen at -20 ^◦^C until protein purification.

##### PDZ3 and synthetic homologs purification

1. Cell pellets were thawed and resuspended in Binding Buffer (20 mmol/L Tris, 500 mmol/L sodium chloride, 10 mmol/L imidazole, pH 7.4) to approximately 0.2 g pellet per mL of Binding Buffer.
2. Resuspended cells were lysed using a 0.125 inch probe sonicator (Fisherbrand Model 120, Thermo Fisher Scientific) for 6 min at 50 % amplitude in cycles (20 s on, 40 s off).
3. Lysates were clarified by centrifugation at 15000 rpm, 4 ^◦^C, for 30 min.
4. Gravity flow columns—one for each purification—were prepared by loading 0.5 mL of nickel-charged sepharose Fast Flow 6 (Cytiva) and washing the sepharose with 3 column volumes (CV) of Binding Buffer.
5. Clarified lysate was flowed through the prepared gravity flow column.
6. Bound protein was washed with 10 CV of Binding Buffer.
7. Bound protein was washed with 6 CV of Wash Buffer (20 mmol/L Tris, 500 mmol/L sodium chloride, 50 mmol/L imidazole, pH 7.4).
8. Bound protein was eluted with 3 CV of Elution Buffer (20 mmol/L Tris, 500 mmol/L sodium chloride, 250 mmol/L imidazole, pH 7.4).
9. Eluted protein was dialyzed at least three times to 1× PBS (137 mmol/L sodium chloride, 2.7 mmol/L potassium chloride, 10 mmol/L sodium phosphate dibasic, 1.8 mmol/L potassium phosphate monobasic pH 7.4) using 2 or 3.5 kDa Slide-A-Lyzer Dialysis cassettes (Thermo Fisher Scientific).
10. Purity of the proteins was evaluated by sodium dodecyl sulfate–polyacrylamide gel electrophoresis (SDS-PAGE) using (4 to 20) % Mini-PROTEAN TGX Precast Protein Gels (Bio-Rad Laboratories). Gels were stained with Coomassie Stain Solution (1.25 g/L Coomassie R250, 50 % methanol, 10 % acetic acid, in water) and destained with De-stain Solution (10 % ethanol, 5 % acetic acid, in water) (fig. S5).

We attempted to purify Homolog 70, but the purification was challenging, even without the DHFR fragment. The synthetic homolog likely requires a significantly modified protocol, due to its low structural and sequence similarity to wildtype.

##### Biolayer interferometry

After dialysis, purified proteins were centrifuged at 21000×*g*, 4 ^◦^C, for 5 min in a chilled tabletop centrifuge to remove any protein that may have crashed out during dialysis. Protein concentration was determined by measuring absorbance at 280 nm (A280) on a Nanodrop spectrophotometer (Thermo Fisher Scientific), using extinction coefficients estimated using the Swiss Institute of Bioinformatics ProtParam tool (*42*, *43*). The highest concentration of each protein sample was prepared with 1× PBS, used for final dialysis, to contain 0.1 % bovine serum albumin (BSA) and 0.05 % Tween-20 to reduce non-specific interactions. Purified proteins were further diluted with Dilution Buffer (0.1 % BSA, 0.05 % Tween-20 in 1× PBS used for final dialysis). After initial tests with serial dilutions of each purified protein, 12.5 *µ*mol/L of protein was used to compare binding activity to each peptide. BLI measurements were performed using the Octet Red96e system (Sartorius) with streptavidin (SA) Biosensors (Sartorius) in 96 well format, measuring 8 wells at a time. Biosensors were hydrated in 200 *µ*L of Dilution Buffer for 10 min at room temperature before loading onto the system. For each measurement, black 96 well plate (Greiner Bio-One) was prepared to have 200 *µ*L/well of each component in successive test columns. The measurements were taken in following steps in the instrument:

1. The peptides (*>*95 % purity by HPLC, Biomatik) used for the BL1 assay were CRIPT (biotin-Ahx-TKNYKQTSV), Syn15 (biotin-Ahx-VSASHDTIV), and Nat18 (biotin-Ahx-IAESLESLV). These were first loaded onto the biosensor tips at 5 *µ*mol/L in the first test column of the 96 well plate.
2. Unbound peptides were washed off by incubating the biosensor tips in Dilution Buffer in the second column of the 96 well plate and obtain a new baseline signal.
3. The binding of the proteins was measured by incubating the biosensor tips in the third column of the 96 well plate with protein samples.
4. The dissociation of the proteins was measured by incubating the biosensor tips in the fourth column of the 96 well plate with Dilution Buffer. The tips were moved along the wells by the instrument.

All measurements were taken at 25 ^◦^C with shaking at 1000 rpm in triplicate (n = 3). The average baseline signal from Step 2 was used to subtract background signal from binding and dissociation response (Steps 3 and 4, respectively) using instrument software Data Analysis HT (Version 12).

#### URA3

##### DNA constructs

Wildtype URA3, controls (R235A and Q215A), and synthetic homologs were inserted to vector pynSPI01 by Twist Bioscience (fig. S14). psynSPI01 is galactose-inducible vector derived from pCTcon2 with yeast-surface display components removed (*44*).

##### General yeast plasmid transformation

Plasmids were transformed into *S. cerevisiae* strain SY992 (MAT*α*/ura3Δ0/leu2Δ0/his3Δ1/lys2Δ0/trp1-63/ade2Δ0) using a modified lithium acetate protocol (*28*). Briefly, SY992 cells were grown to log phase in Yeast Peptone Dextrose (YPD), harvested, and then washed with TE buffer (10 mmol/L Tris, 1 mmol/L ethylenediaminetetraacetic acid (EDTA)) and lithium acetate mix (100 mmol/L lithium acetate, 10 mmol/L Tris, 1 mmol/L EDTA). The cells were resuspended in final lithium acetate mix at a ratio of 200 *µ*L per 5 mL of source YPD culture (*i.e.*, 25 mL of culture yielded 1 mL of resuspended cells). Transformations were then performed in low throughput using 1.5 mL microcentrifuge tubes or high throughput using 96 well flat bottom polystyrene plates (Falcon). After transformation, cryogenic stocks were prepared for each plasmid with overnight cultures from three individual colonies grown in SD-Trp (6.74 g/L yeast nitrogen base with ammonium sulfate, 0.72 g/L CSM-Trp powder, 20 g/L dextrose, pH 5.8) at a final glycerol concentration of 20 %.

### Low-throughput yeast plasmid transformation

1. For each unique transformation, a mixture was prepared in a 1.5 mL microcentrifuge tube: 350 *µ*L PEG-3350 mix (40 % (w/v) PEG-3350 in Nanopure water, 100 mmol/L lithium acetate, 10 mmol/L Tris, 1 mmol/L EDTA), 10 *µ*L boiled salmon sperm ssDNA (Thermo Fisher Scientific), and 200 ng plasmid DNA.
2. To each mixture, 50 *µ*L of transformation prepared cells were added.
3. The mixture was vortexed and incubated at room temperature for 30 min.
4. 24 *µ*L of DMSO was added to each mixture, the mixtures briefly vortexed, then incubated at 42 ^◦^C for 15 min.
5. Cells were pelleted by centrifugation at 8000×*g*, room temperature, for 1 min and the transformation mix supernatant removed by pipetting.
6. Cells were then resuspended in 200 *µ*L YPD and 100 *µ*L of the resuspension was plated on SD-Trp agar using glass beads.
7. Plates were incubated at 30 ^◦^C.

### High-throughput yeast plasmid transformation

1. A mixture of 175 *µ*L PEG-3350 mix and 5 *µ*L boiled salmon sperm ssDNA per transformation was prepared as a master mix and distributed into a 96 well polystyrene plate.
2. 100 ng plasmid DNA (1 *µ*L) was added into each well with master mix.
3. 50 *µ*L transformation prepared cells were added to each well and vigorously mixed by pipetting.
4. The plate was incubated at room temperature for 30 min.
5. 12 *µ*L of DMSO was added to each well and vigorously mixed by pipetting.
6. The plate was incubated at 42 ^◦^C for 15 min.
7. Cells were pelleted by centrifugation at 5000×*g*, room temperature, for 1 min and the transformation mix supernatant removed by pipetting.
8. Cells were resuspended in 100 *µ*L YPD and 50 *µ*L of the resuspension was plated on SD-Trp agar using glass beads.
9. Plates were incubated at 30 ^◦^C.

#### URA3 cellular assay

Four different media conditions were used to set up and run the growth-based assay:

- SD-Trp to initially grow cells containing plasmid to set up for the assay.
- SG-Trp-Ura for the main assay condition, where cell growth was dependent on expression of active URA3 induced by galactose (6.74 g/L yeast nitrogen base with ammonium sulfate, 0.72 g/L CSM-Trp-Ura powder, 20 g/L galactose, pH 5.5).
- SD-Trp-Ura for the negative control condition, where URA3 was not expressed (6.74 g/L yeast nitrogen base with ammonium sulfate, 0.72 g/L CSM-Trp-Ura powder, 20 g/L dextrose, pH 5.5).
- SG-Trp for the positive control condition, where cells grew independent of URA3 expression from plasmid (6.74 g/L yeast nitrogen base with ammonium sulfate, 0.72 g/L CSM-Trp powder, 20 g/L galactose, pH 5.8).

Overnight cultures were started from glycerol scrapings in 1.5 mL SD-Trp medium in 14 mL snapcap culture tubes. Cells were grown at 30 ^◦^C, 300 rpm, for 24 h. Aliquots of SD-Trp, SD-Trp-Ura, SG-Trp, and SG-Trp-Ura media were loaded on the automation system along with the overnight cultures.

The following processes were carried out with an automated liquid handling platform in sequential order:

1. Growth Plate 1 preparation: 450 *µ*L of SD-Trp was mixed with 50 *µ*L of overnight cultures (1:10 dilution of cells).
2. Growth Plate 1 measurement: Growth Plate 1 was sealed with a breathable membrane and moved to a Biotek Epoch2 reader (Agilent Technology). Plates were incubated at 30 ^◦^C, with shaking, and OD_600_ measured every 10 min for 12 h.
3. Growth Plate 2 preparation: 10 *µ*L of cells from Plate 1 were mixed with 490 *µ*L of one of following media: SD-Trp-Ura, SG-Trp-Ura, or SG-Trp.
4. Growth Plate 2 measurement: Growth Plate 2 was sealed with a breathable membrane and moved to a Biotek Epoch2 reader (Agilent Technology). Plates were incubated at 30 ^◦^C, without shaking, and OD_600_ measured every 10 min for 48 h.

*Metrics:* Normalized activity score was recorded as:

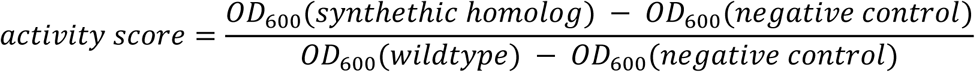

at 24 h of growth of Plate 2. The negative control was defined as cells carrying the URA3 Q215A sequence (negligible catalytic activity) grown in SD-Trp-Ura media used to maintain plasmid but not induce protein expression. Any synthetic homologs with growth significantly exceeding the negative control as determined by a Welch’s one-tailed t-test (*p<*0.05) were labeled active.

#### T7 RNAP

##### DNA constructs

The promoter J23119 used for T7 RNAP plasmid templates was obtained from the Anderson promoter collection in the Standard Registry of Biological Parts. Plasmid pJL1 expressing superfolder GFP (sfGFP), a gift from Michael Jewett (Addgene plasmid 69496) (*45*), was used as the reporter plasmid. sfGFP was selected as the fluorescent output in cell-free reactions because it matures rapidly and produces a strong signal. For cell-based assays, the mScarlet-I3 gene was inserted into pJL1 to replace sfGFP via Gibson assembly (E2611L, New England Biolabs). Plasmids containing T7 RNAP and a mutated, nonfunctional promoter were ordered from Twist Bioscience using their “pTwist Kan Medium Copy” cloning vector (fig. S14). *E. coli* DH10*β* and LB medium were used for all cell growth during the cloning steps. Kan (33 *µ*g/mL) and chloramphenicol (25 *µ*g/mL) were used for appropriate antibiotic selection.

##### General setup

T7 RNAP and its reporter were expressed from two separate cassettes (fig. S14). The first cassette expressed the T7 RNAP synthetic homolog and a fluorogenic Pepper aptamer from an *E. coli* promoter (J23119) (*46*). The Pepper aptamer was intended for transcript level monitoring. The T7 cassette included insulators for transcription and translation (RiboJ and BCD, respectively) to account for sequence-specific variations in expression (*39*, *40*). The reporter cassette expressed a fluorescent protein (FP) from the T7 promoter.

### Cell-free expression (CFE) assays

#### Bacterial extract preparation

Extracts for cell-free reactions were prepared as previously described (*31*). Briefly:

1. *E. coli* BL21 Δ*recBCD* cells (Addgene product 176581, a gift from Chase Beisel & Jean-Loup Faulon) (*47*) were grown in 2× YTP medium (5 g/L sodium chloride, 10 g/L yeast extract, 16 g/L tryptone, 40 mmol/L potassium phosphate dibasic, 22 mmol/L potassium phosphate monobasic) at 37 ^◦^C, 250 rpm.
2. Upon reaching mid-exponential phase (OD_600_ ∼1.6-1.7), cells were harvested by centrifugation at 3000×*g*, 4 ^◦^C, for 15 min, and then washed three times with S30 buffer (14 mmol/L magnesium acetate, 60 mmol/L potassium acetate, 10 mmol/L Tris-acetate, pH adjusted to 8.2 with potassium hydroxide, and 2 mmol/L dithiothreitol added immediately before use).
3. Cells were resuspended in 1 mL of S30 buffer per g of cells, and this suspension was transferred to a 1.5-mL microcentrifuge tube in 1 mL aliquots.
4. The cell resuspension was sonicated (Q125 sonicator, QSonica) on ice at 20 kHz frequency, and 50 % amplitude in cycles (10 s on, 10 s off) until the cumulative sonication energy input reached 250 ± 10 J.
5. Immediately after lysis, 3 mmol/L dithiothreitol was added to each tube.
6. Lysed cells were centrifuged at 12000×*g*, 4 ^◦^C, for 15 min.
7. The supernatant was consolidated and placed in an incubator shaker at 37 ^◦^C, 250 rpm, for 80 min.
8. The samples were centrifuged at 12000×*g*, 4 ^◦^C, for 15 min, after which the supernatant (*i.e.*, the final extract to be used in cell-free reactions) was consolidated, mixed by inversion, and divided into 250 *µ*L aliquots in 1.5-mL microcentrifuge tubes. The aliquots were stored at -80 ^◦^C until use.

#### Linear T7 RNAP templates

Linear templates were prepared by PCR and subsequently purified by spin column extraction. The 5’ PCR primer restores the J23119 promoter region in the T7 RNAP constructs, which were originally synthesized and cloned with nonfunctional promoter to mitigate cytotoxicity concerns in DNA synthesis and purification.

#### Plasmid T7 RNAP templates

A subset of T7 RNAP synthetic homologs were successfully cloned into a plasmid with a functional J23119 promoter. These plasmids were tested in a cell-free assay to circumvent the variability observed across PCR preparations.

#### Cell-free reaction assays

Cell-free reactions were prepared as previously described (*31*). Reactions for all experiments contained 10 mmol/L magnesium glutamate, 10 mmol/L ammonium glutamate, 133 mmol/L potassium glutamate, 1.2 mmol/L ATP, 0.85 mmol/L GTP, 0.85 mmol/L CTP, 0.85 mmol/L UTP, 0.034 mg/mL folinic acid, 0.171 mg/mL tRNA from *E. coli* MRE 600, 0.33 mmol/L nicotinamide adenine dinucleotide (NAD), 0.2667 mmol/L coenzyme A (CoA), 4 mmol/L sodium oxalate, 1 mmol/L putrescine, 1.5 mmol/L spermidine, 57 mmol/L HEPES, 5 *µ*mol/L HBC620 dye, 2 mmol/L each of the twenty standard amino acids, 0.03 mol/L PEP, 27 % (v/v) cell extract, and the nucleic acid template concentration specified for each experiment.

Reactions were run in 10 *µ*L volumes in clear 384 well plates (Greiner Bio-One) in a BioTek Synergy Neo2 reader (Agilent Technology) at 37 ^◦^C. A clear adhesive film (Thermo Fisher Scientific) was used to cover the plate and prevent evaporation. Excitation and emission wavelengths were, respectively, 485 nm and 510 nm for sfGFP, and 577 nm and 620 nm for the HBC620 fluorophore. In reactions with DNA templates (both linear and plasmid), 15 nmol/L of T7 RNAP-expressing template and 5 nmol/L of fluorescent protein-expressing template were added. In reactions with RNA templates, 100 nmol/L of T7 RNAP-Pepper RNA were added to ensure that the same amount of T7 RNAP mRNA transcripts were present, which could not be reliably monitored with the Pepper aptamer.

#### In vitro transcription

*In vitro* transcription reactions to generate T7 RNAP-expressing mRNAs contained 25 mmol/L linear DNA template, 2 mmol/L each of ATP, GTP, CTP and UTP, 15 U/*µ*L *E. coli* RNA polymerase (M0551S, New England Biolabs), and the appropriate buffer provided by the polymerase manufacturer. Reactions were run in 150 *µ*L volumes at 37 ^◦^C for 2 h in a thermocycler. Each reaction was then treated with 50 mmol/L EDTA and 0.027 U/*µ*L DNase I at 37 ^◦^C for 15 min prior to purification and elution in 30 *µ*L nuclease-free water.

##### Cell-based Assay

The assay used *E. coli* BL21 as the chassis and the two cassettes described in “General Setup” in a plasmid format. mScarlet-I3 was chosen as the fluorescent protein reporter for this assay due to lower cell autofluorescence within the red spectrum. *E. coli* BL21 cells (C2530H, New England Biolabs) were co-transformed with two plasmids, one expressing T7 RNAP from a constitutive *E. coli* promoter (J23119) and a second expressing mScarlet-I3 from a constitutive T7 promoter. Three colonies from each transformation plate were inoculated overnight in LB medium supplemented with antibiotics to maintain the plasmids. 6 *µ*L of the overnight culture were then added to 294 *µ*L of LB (with antibiotics), and the samples were placed in a BioTek Synergy Neo2 reader (Agilent Technology) at 37 ^◦^C for 6 h, after which they were transferred to a flow cytometer for analysis. For flow cytometry, 5 *µ*L of cell culture were mixed with 195 *µ*L of PBS and 170 *µ*g/mL of chloramphenicol. Cells harboring the mScarlet-I3 plasmid only were used as a negative control. BL21 Star (DE3) cells containing mScarlet-I3 plasmid and expressing wildtype T7 RNAP were used as a positive control due to a cytotoxicity issue in BL21 cells. All reactions were run in a clear, flatbottomed 96 well plate. Excitation and emission wavelengths for mScarlet-I3 were 568 nm and 592 nm, respectively.

#### Metrics

For both CFE and cell-based assays, normalized activity score was calculated as:

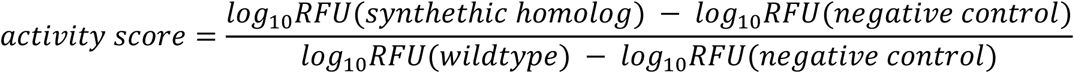

at 6 h post DNA addition.

