## Supplemental file for "Experimental evaluation of AI-driven protein design risks using safe biological proxies"

**The PDF file includes:**

Figs. S1 to S14
Tables S1 to S2

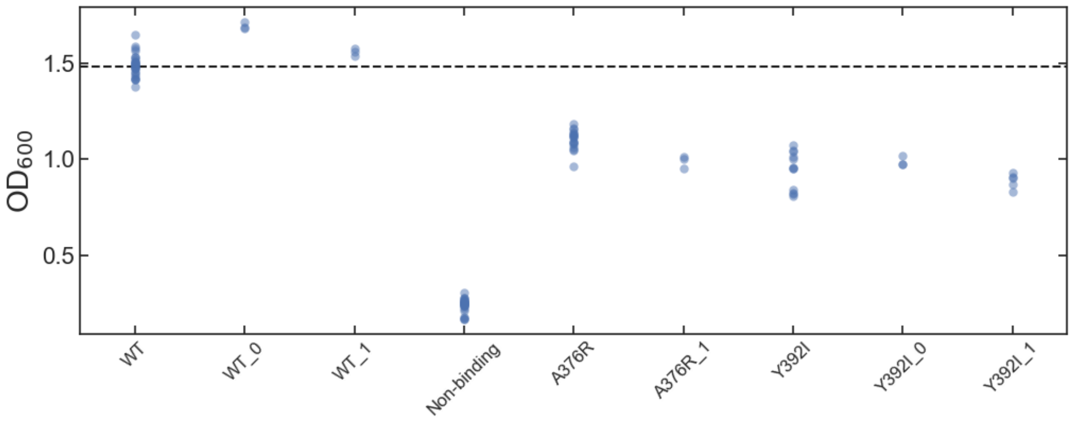


**Fig. S1.**

**PDZ3-CRIPT DHFR complementation assays with controls.** Endpoint optical density measured at 600 nm ($OD_{600}$) for each sample is displayed, with each point representing a measurement. The non-binding control contains PDZ3 and DHFR fragments but not CRIPT. Control sequence variants (A376R and Y392I) are named in reference to the wildtype sequence. Subscripts indicate different codon optimization schemes. The dotted line is the mean of the wildtype PDZ3 control (WT) (n = 28).

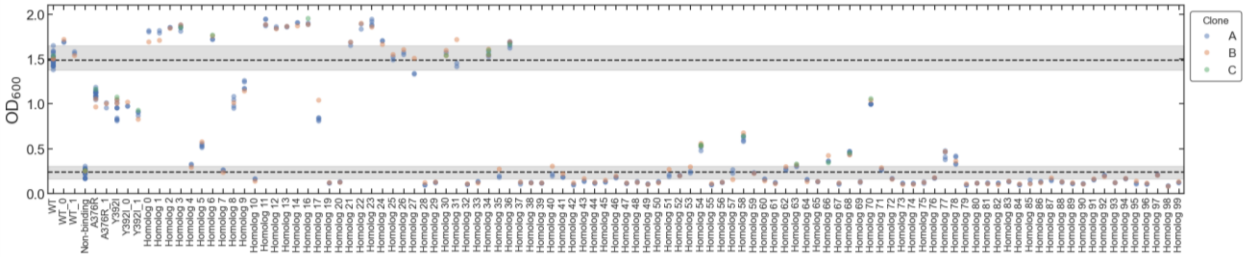


**Fig. S2.**

**PDZ3-CRIPT DHFR complementation assays with synthetic homologs.** Endpoint optical density measured at 600 nm ($OD_{600}$) for each sample is displayed, with each point representing a measurement and each color representing a transformant clone. The non-binding control contains PDZ3 and DHFR fragments but not CRIPT. Control sequence variants (A376R and Y392I) are named in reference to the wildtype sequence. Subscripts indicate different codon optimization schemes. The dotted lines indicate the mean values of the non-binding control (PDZ3 and DHFR fragments only) (n = 28) and wildtype PDZ3 control (WT) (n = 28). Grey shaded regions around the mean values indicate the minimum and maximum range of $OD_{600}$ measured for the control samples.

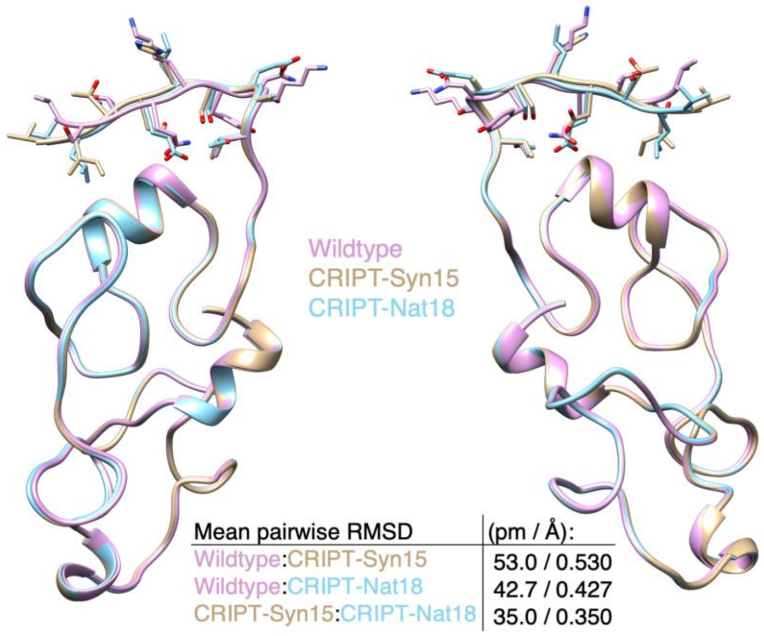


**Fig. S3.**

**Alignment of CRIPT with modified C-terminal peptide sequences.** Models were generated with AlphaFold 3 and aligned using UCSF Chimera's MatchMaker tool to determine whether C-terminus modification impacts the overall CRIPT geometry (*1*, *2*). The root mean square deviation (RMSD) is reported in picometers (pm) and Angstroms (Å). It should be noted that due to the reducing environment in the cytosol of *E. coli*, the structures may fold differently than predicted.

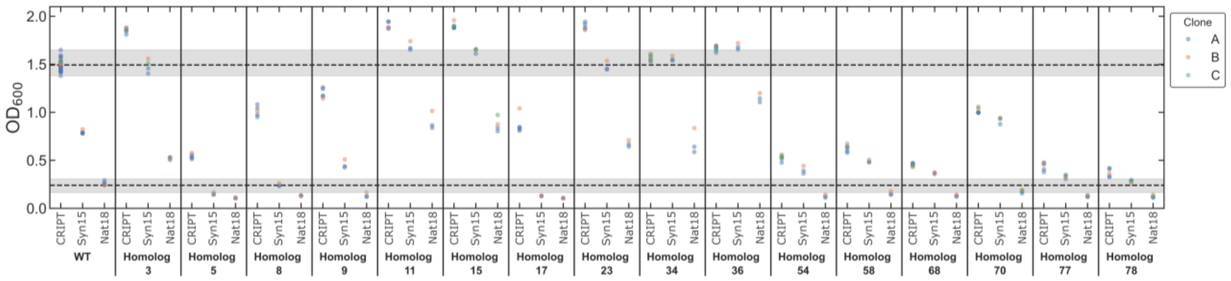


**Fig. S4.**

**Results of PDZ3-CRIPT DHFR complementation assays with modified CRIPT C-terminus for a subset of PDZ3 synthetic homologs.** Endpoint optical density measured at 600 nm ($OD_{600}$) for each sample is displayed, with each point representing a measurement and each color representing a transformant clone. The x-axis label indicates whether the peptide at C-terminus of CRIPT was wildtype (CRIPT) or control peptides (Syn15 or Nat18). The dotted lines indicate the mean values of the non-binding control (PDZ3 and DHFR fragments only) (n = 28) and wildtype PDZ3 control (WT) (n = 28). Grey shaded regions around the mean values indicate the minimum and maximum range of $OD_{600}$ measured for the control samples.

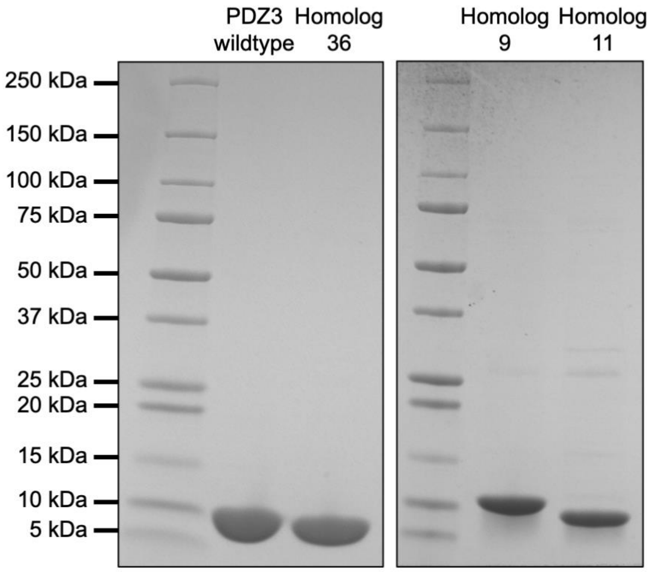


**Fig. S5.**

**Representative SDS-PAGE gels of purified PDZ3 synthetic homologs.** The estimated molecular mass of the 6×histidine-tagged wildtype PDZ3 domain is 10.6 kDa, calculated with the Swiss Institute of Bioinformatics ProtParam tool (*3*, *4*). Brightness and contrast adjustments have been applied to the entire gel images for each gel.

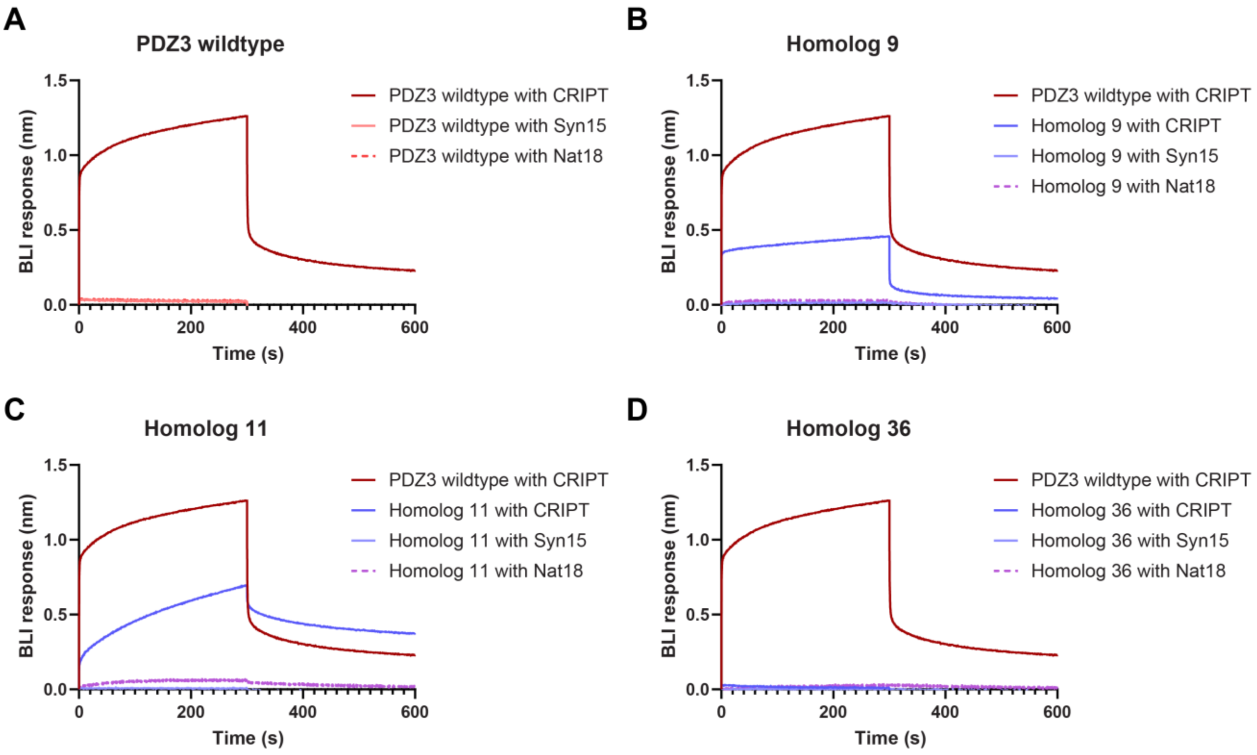


**Fig. S6.**

**Binding response of purified PDZ3 proteins against CRIPT, Syn15, and Nat18 peptides.** Representative data of BLI response of (**A**) PDZ3 wildtype, (**B**) Homolog 9, (**C**) Homolog 11, and (**D**) Homolog 36. After immobilization of the corresponding peptide onto the streptavidin biosensor tips, the tips were incubated with 12.5 $\mu$mol/L of purified protein to obtain binding curves and then in Dilution Buffer to obtain dissociation curves (n = 3). The binding response of PDZ3 wildtype with CRIPT is included in all plots to facilitate comparison.

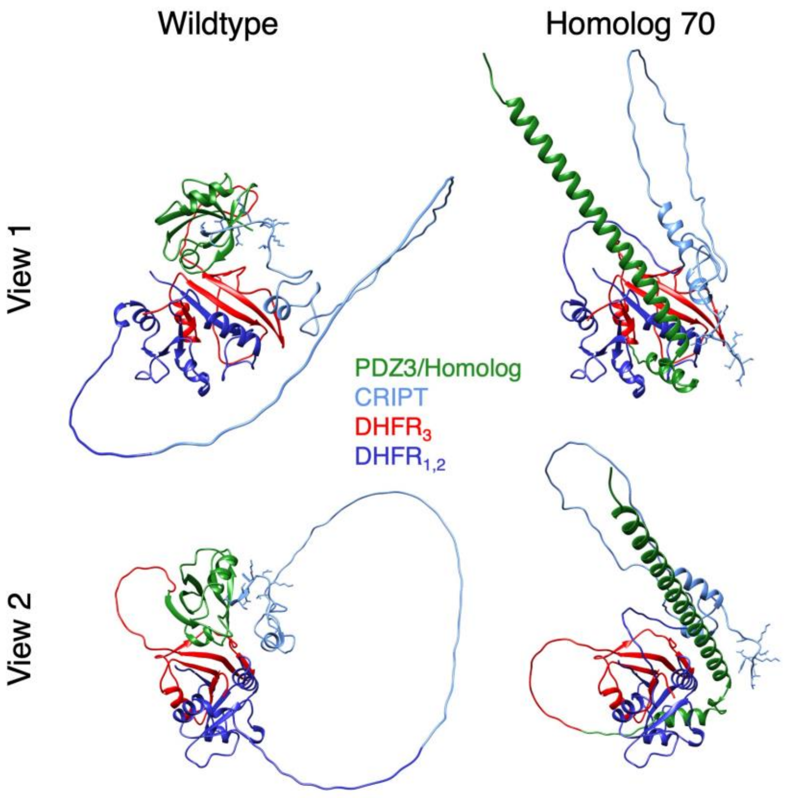


**Fig. S7.**

**Structural modeling of interactions of DHFR-CRIPT with PDZ3 and Homolog 70.** AIPD-generated Homolog 70 appeared to have activity above the threshold in the primary cellular assay, despite very low sequence identity to canonical PDZ3 (2.38 %). An AlphaFold3 prediction showed a dramatically different helix-turn-helix conformation of Homolog 70 and its interaction with CRIPT compared to wildtype (*1*). It should be noted that the reducing environment of the *E. coli* cytosol is not accounted for in the prediction. Regardless, this result highlights unforeseen, confounding effects of AI-generated sequences that could occur in the transition from *in silico* modeling to experimental validation in cells.

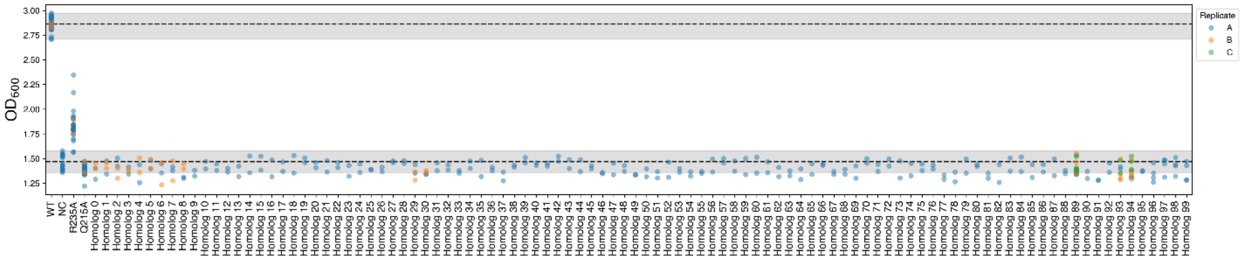


**Fig. S8.**

**URA3 cellular assay with synthetic homologs.** Optical density measured at 600 nm ($OD_{600}$) for each sample at 24 h of growth displayed, with each point representing a measurement and each color representing a transformant clone. Control sequence variants (Q215A and R235A) are named in reference to the wildtype sequence. The negative control (NC) was the severely attenuated variant Q215A in negative control medium SD-Trp-Ura (*5*). The dotted lines indicate the mean values of the NC (n = 18) and wildtype URA3 control (WT) induced in SG-Trp-Ura medium (n = 26). All synthetic homologs were induced in SG-Trp-Ura medium. Grey shaded regions around the mean values indicate the minimum and maximum range of $OD_{600}$ measured for the control samples.

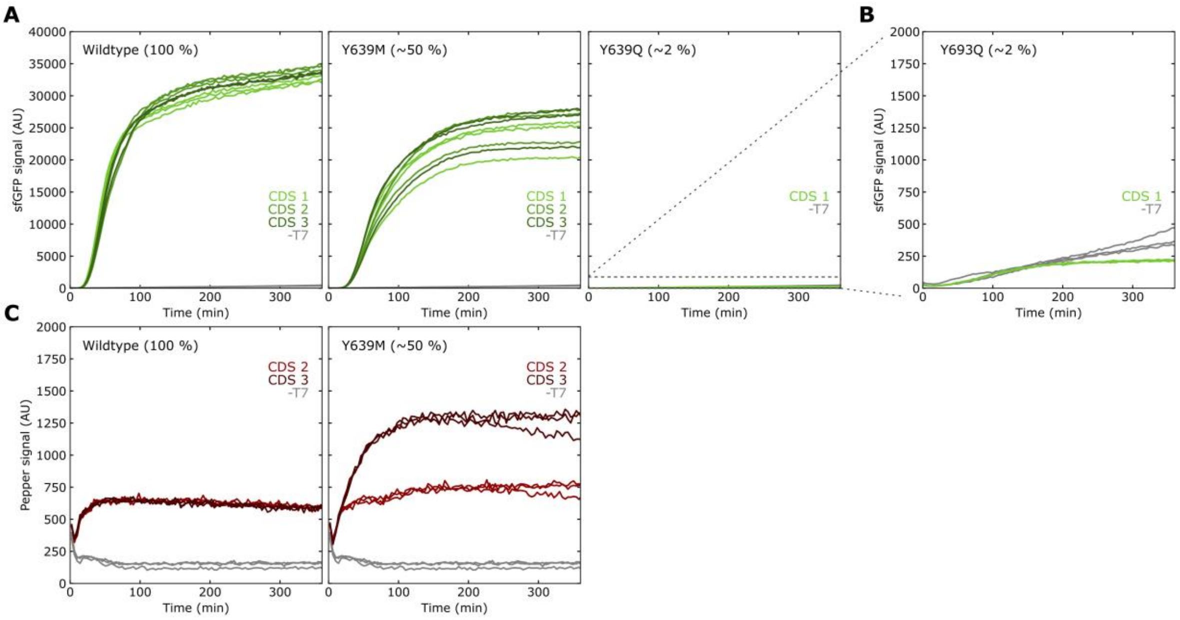


**Fig. S9.**

**Validation and bounds of CFE assay to measure T7 RNAP activity.** (**A**) sfGFP expression induced by *in situ* production of T7 RNAP controls in our CFE system. The Y639M and Y639Q controls have point mutations reported to reduce T7 RNAP activity to 50 % and 2 % of wildtype, respectively (*6*). Three wildtype and Y639M codon schemes (CDSs) were tested to determine if codon choice would substantially influence our measurements (n = 3). (**B**) Replotting of the Y639Q data on narrower y-axis. The CFE assay was unable to distinguish this control variant from background (-T7 control: T7 plasmid absent, fluorescent reporter plasmid only). Individual traces represent separate technical replicates for each construct (n = 3). (**C**) Pepper aptamer signal for the T7 RNAP transcripts of a subset of the variants tested in (A) (n = 3). Different CDSs did not result in substantial difference in sfGFP expression, the reporter for T7 RNAP activity. The Pepper signal did differ substantially for one CDS, suggesting that Pepper was sensitive to sequence context and is not a reliable signal for normalizing transcript levels across synthetic homologs.

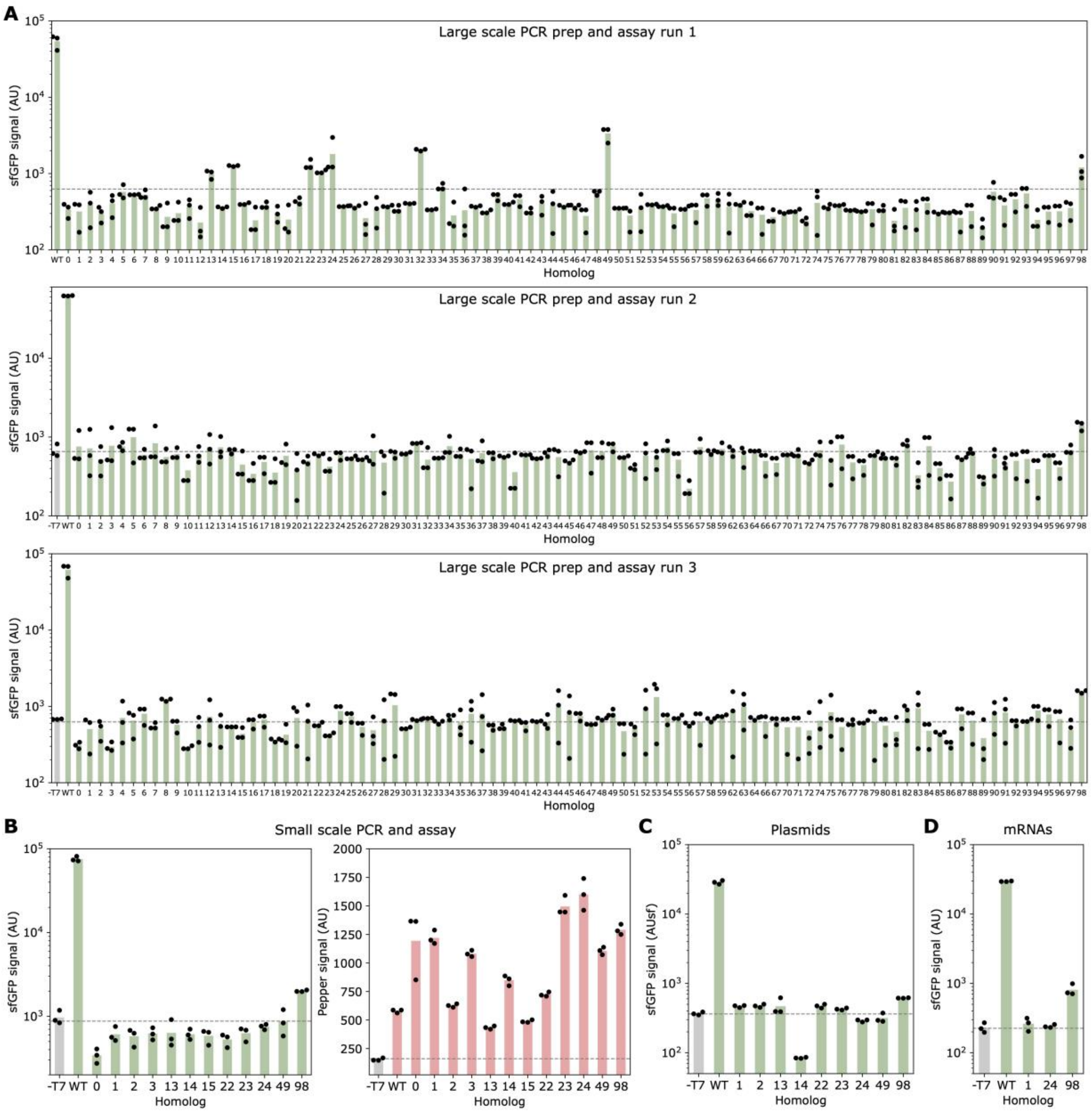


**Fig. S10.**

**Cell-free expression assay results with T7 RNAP synthetic homologs.** (**A**) Two independent replicates of large-scale measurements of 93 T7 RNAP synthetic homologs. PCR, PCR purification, and assay preparations were conducted independently for each replicate in a 96 well plate format with multichannel pipettes. Each data point indicates a technical replicate performed in the same 384 well plate (n = 3). (**B**) To rule out experimental variation due to manual pipetting, a subset of synthetic homologs were retested individually at a smaller scale. Each data point indicates a different PCR and PCR purification for the same sample, measured in triplicate on the same plate (n = 3). sfGFP signal (left) and the corresponding Pepper aptamer signal (right) are plotted for each T7 RNAP synthetic homolog. Pepper signal differs substantially across synthetic homologs and does not appear to correlate with sfGFP expression levels, as observed in fig. S9C. (**C**) To address concerns of the PCRs introducing mutations in the constructs, as well as undesired dsDNA degradation of the linear PCR templates in our CFE lysate, selected plasmid templates were prepared from expansion in DH10$\beta$ cells. Each data point indicates a technical replicate consisting of plasmids from a single extraction (n = 3). (**D**) To rule out variations in transcription across DNA templates, *in vitro* transcription was conducted for a select set of synthetic homologs. Fixed amount of purified and quantified mRNA was then added to CFE (n = 3). All results were quantified after 6 hr. Dashed gray lines indicate the background measurement from a -T7 control (T7 plasmid absent, fluorescent reporter plasmid only).

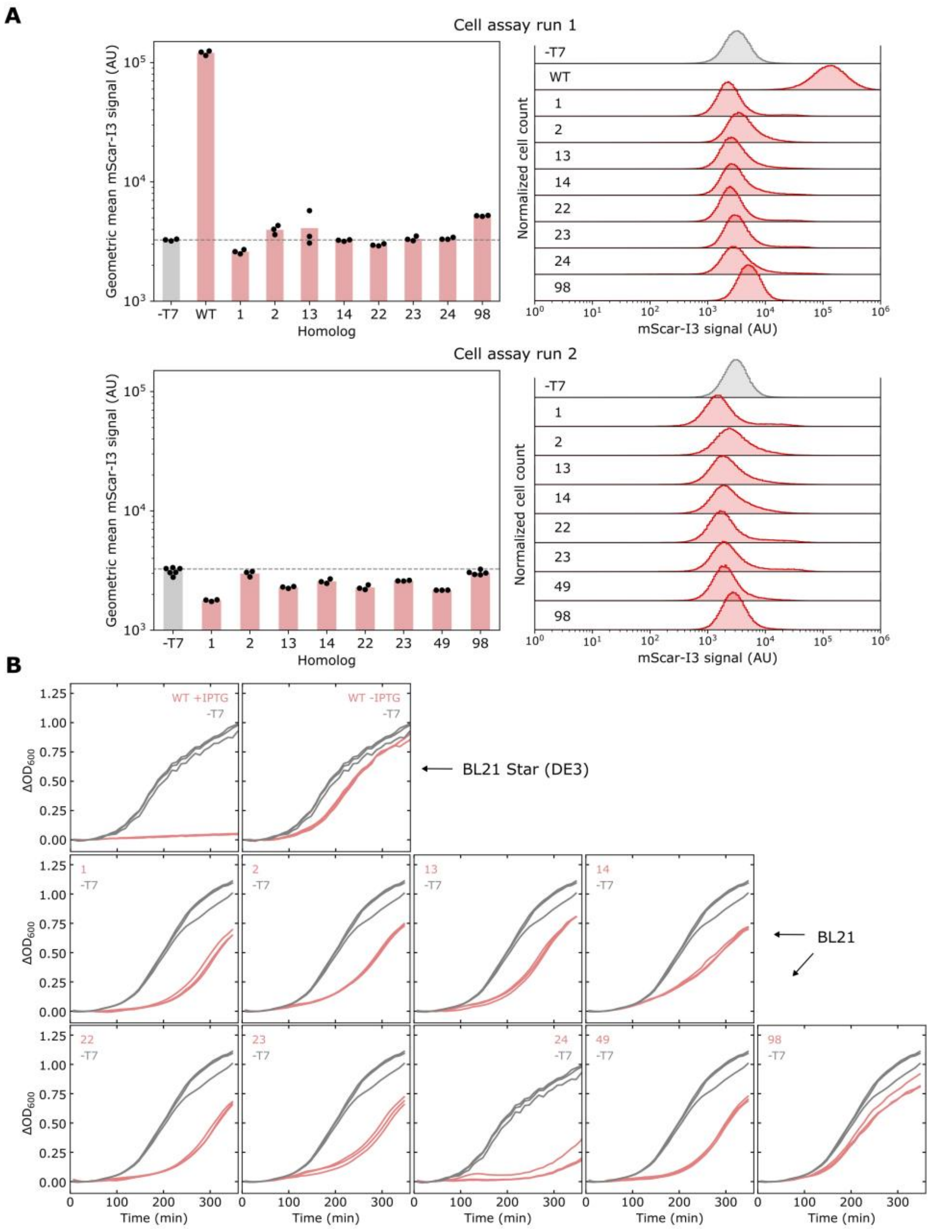


**Fig. S11.**

***E. coli*** **assay results with T7 RNAP synthetic homologs.** (**A**) Flow cytometry results for select T7 RNAP synthetic homologs. Left: geometric means of the distributions. Right: fluorescent cell distributions. Each data point indicates a sample originating from a different transformant ($n \geq 3$). Dashed gray lines indicate the background level from the -T7 control (T7 plasmid absent, Scarlett-I3 reporter plasmid only). The wildtype (WT) control contained inducible T7 RNAP in its genome and the reporter plasmid. (**B**) Growth curves of strains containing T7 RNAP synthetic homolog and fluorescent reporter plasmids. Growth curves were obtained by measuring absorbance at 600 nm over time and subtracting the initial absorbance value at time = 0 min ($\Delta OD_{600}$). Individual traces indicate biological replicates originating from different transformants ($n \geq 3$). Induction of T7 RNAP expression with 100 µmol/L of IPTG (WT + IPTG) was toxic, so the wildtype control shown in (**A**) was not induced (basal expression of T7 RNAP in the strain).

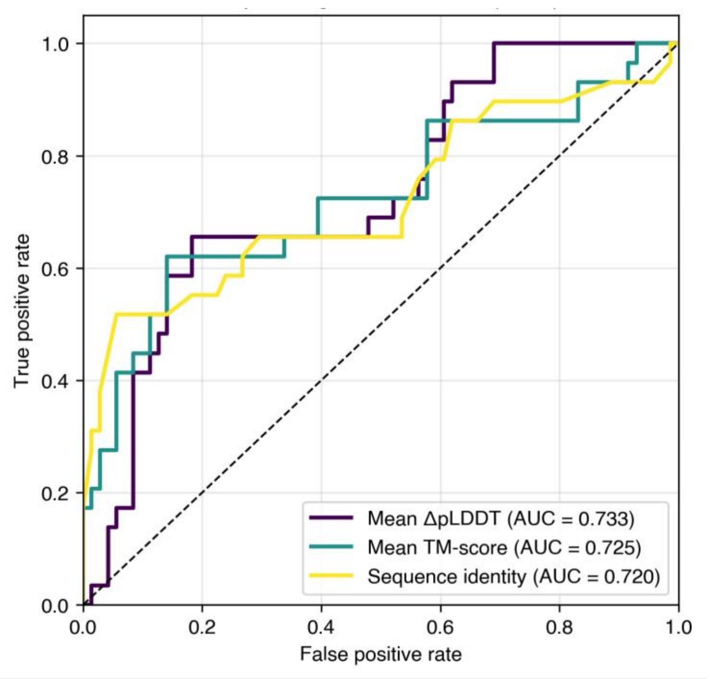


**Fig. S12.**

**Receiver Operating Characteristic curves for PDZ3 AIPD metrics used to predict activity label.** Mean ΔpLDDT (purple, AUC = 0.733), mean TM-score (aqua, AUC = 0.725), and sequence identity (yellow, AUC = 0.720) all show similar predictive power and strong covariance (mean Spearman's $\rho = 0.91$). The interdependence of sequence and structural metrics for PDZ3 may explain AIPD and BSS success, which was only achieved for this protein target. True positive rate (sensitivity) is plotted against false positive rate (1 − specificity) across varying classification thresholds. All metrics substantially outperform random classification (dashed diagonal line), indicating that the AIPD pipeline does have a basic level of activity prediction capability.

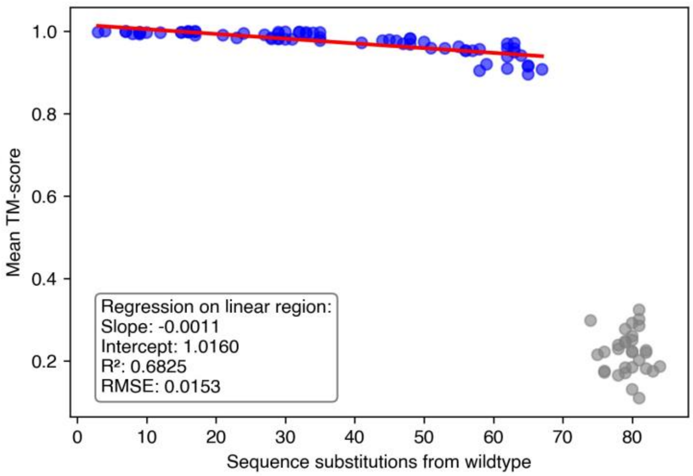


**Fig. S13.**

**Linear regression analysis of sequence identity versus TM-score for PDZ3 synthetic homologs.** The x-axis counts the number of amino acid substitutions for each synthetic homolog compared to wildtype PDZ3, and the y-axis shows the mean TM-score calculated between wildtype PDZ3 and 200 models generated by OpenFold from unique seeds. Each point represents a synthetic homolog ($n = 100$). As sequence identity to wildtype decreases, mean TM-score decreases linearly (blue region) until abruptly decreasing at 74 substitutions (grey circles). This suggests the PDZ3 fold is highly permissive to drastic sequence changes below approximately 70 sequence substitutions.

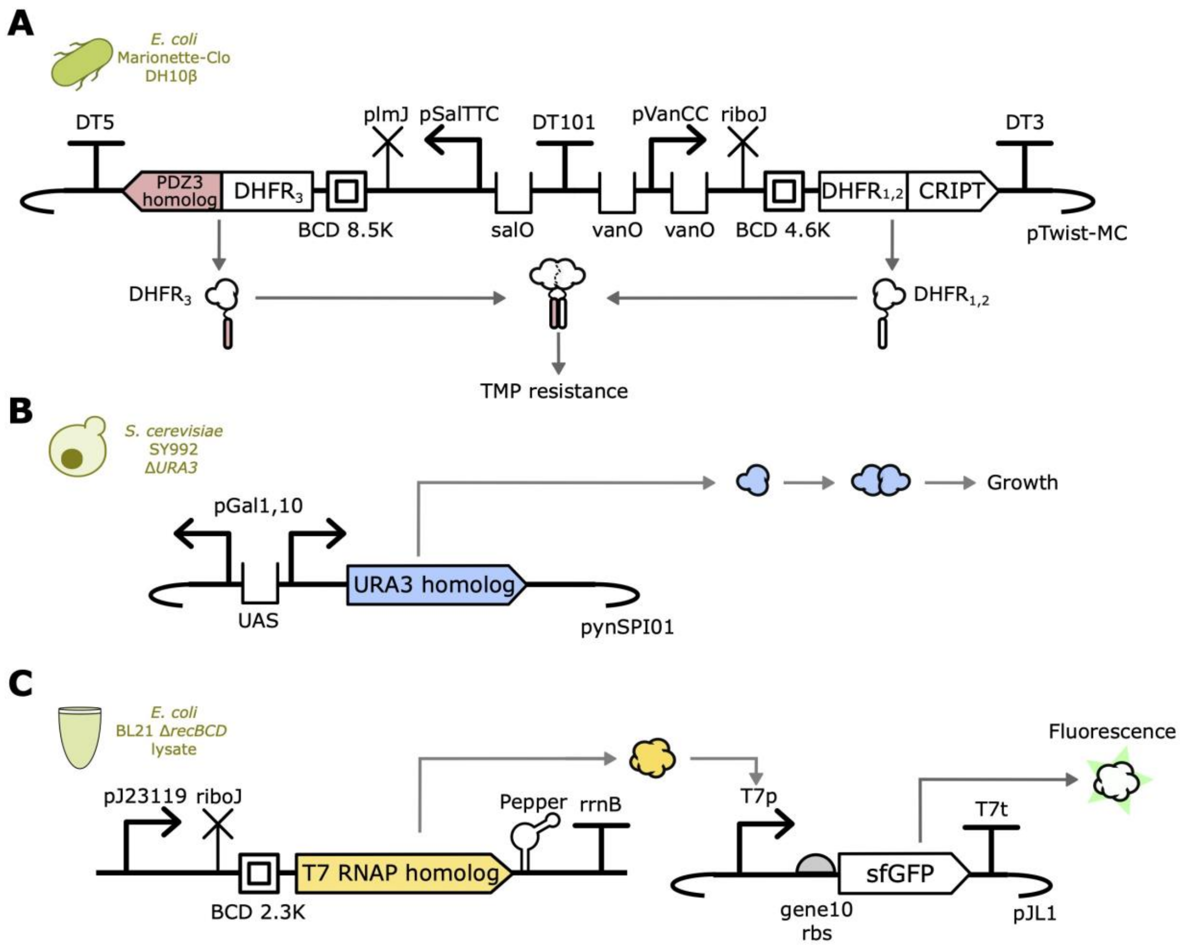


**Fig. S14.**

**Plasmid schematics for primary TEVV assays.** (**A**) For the PDZ3 cellular assay, $DHFR_{1,2}$-CRIPT and $DHFR_3$-PDZ3 synthetic homolog fusions were expressed from a single plasmid containing bicistronic design translational control elements (BCDs) to reduce sequence specific variation in expression. Interaction of wildtype PDZ3 or synthetic homolog with CRIPT brings together the DHFR fragments to reconstitute active enzyme and confer TMP resistance. (**B**) For the URA3 cellular assay, URA3 expression was induced by galactose via the pGal1,10 promoter on a plasmid (pynSPI01). Expression of active URA3 leads to uracil biosynthesis and cell growth in a uracil-free media. **(C)** For T7 RNAP CFE assays, a linear template (left) expressed a T7 RNAP synthetic homolog and the fluorogenic Pepper aptamer from an *E. coli* promoter (J23119). The template included insulators for transcription and translation (riboJ and BCD, respectively) to account for sequence-specific variations in expression. The T7 RNAP synthetic homolog transcribed a superfolder GFP (sfGFP) gene encoded in a plasmid under a T7-specific promoter (right).

**Table S1.**

**Key residue positions held constant during synthetic homolog generation.** The AIPD pipeline is able to accept user-supplied key residue positions to hold constant during synthetic homolog generation. For each protein target, we reviewed structures, literature, and/or a mutational consequence database (for enzymes) to select key wildtype residues (*7*).

| Protein | UniProtID | PDB ID | Constant Position | Reason | Citation |
|---|---|---|---|---|---|
| PDZ3 | P78352 | 3I4W | LEU 323 | Literature evidence of importance | (*8*) |
| PDZ3 | P78352 | 3I4W | GLY 324 | Literature evidence of importance | (*8*) |
| PDZ3 | P78352 | 3I4W | PHE 325 | Literature evidence of importance | (*8*) |
| PDZ3 | P78352 | 3I4W | ILE 327 | Literature evidence of importance | (*8*) |
| PDZ3 | P78352 | 3I4W | VAL 328 | Literature evidence of importance | (*8*) |
| PDZ3 | P78352 | 3I4W | GLY 329 | Literature evidence of importance | (*8*) |
| PDZ3 | P78352 | 3I4W | GLY 330 | Literature evidence of importance | (*8*) |
| PDZ3 | P78352 | 3I4W | ILE 336 | Literature evidence of importance | (*8*) |
| PDZ3 | P78352 | 3I4W | ILE 338 | Literature evidence of importance | (*8*) |
| PDZ3 | P78352 | 3I4W | ILE 341 | Literature evidence of importance | (*8*) |
| PDZ3 | P78352 | 3I4W | ALA 347 | Literature evidence of importance | (*8*) |
| PDZ3 | P78352 | 3I4W | LEU 353 | Literature evidence of importance | (*8*) |
| PDZ3 | P78352 | 3I4W | ILE 359 | Literature evidence of importance | (*8*) |
| PDZ3 | P78352 | 3I4W | VAL 362 | Literature evidence of importance | (*8*) |
| PDZ3 | P78352 | 3I4W | LEU 367 | Literature evidence of importance | (*8*) |
| PDZ3 | P78352 | 3I4W | HIS 372 | Literature evidence of importance | (*8*) |
| PDZ3 | P78352 | 3I4W | ALA 375 | Literature evidence of importance | (*8*) |

| | | | | | |
|---|---|---|---|---|---|
| PDZ3 | P78352 | 3I4W | ALA 376 | Literature evidence of importance | (*8*) |
| PDZ3 | P78352 | 3I4W | LEU 379 | Literature evidence of importance | (*8*) |
| PDZ3 | P78352 | 3I4W | ILE 388 | Literature evidence of importance | (*8*) |
| URA3 | P03962 | 3gdl | ASP 37 | Substrate binding | UniProt entry |
| URA3 | P03962 | 3gdl | LYS 59 | Substrate binding | UniProt entry |
| URA3 | P03962 | 3gdl | THR 60 | Substrate binding | UniProt entry |
| URA3 | P03962 | 3gdl | HIS 61 | Substrate binding | UniProt entry |
| URA3 | P03962 | 3gdl | ASP 91 | Substrate binding | UniProt entry |
| URA3 | P03962 | 3gdl | ARG 92 | Substrate binding | UniProt entry |
| URA3 | P03962 | 3gdl | LYS 93 | Proton donor | UniProt entry |
| URA3 | P03962 | 3gdl | PHE 94 | Substrate binding | UniProt entry |
| URA3 | P03962 | 3gdl | ALA 95 | Substrate binding | UniProt entry |
| URA3 | P03962 | 3gdl | ASP 96 | Substrate binding | UniProt entry |
| URA3 | P03962 | 3gdl | ILE 97 | Substrate binding | UniProt entry |
| URA3 | P03962 | 3gdl | GLY 98 | Substrate binding | UniProt entry |
| URA3 | P03962 | 3gdl | ASN 99 | Substrate binding | UniProt entry |
| URA3 | P03962 | 3gdl | THR 100 | Substrate binding | UniProt entry |
| URA3 | P03962 | 3gdl | SER 154 | Substrate binding | BRENDA (*17*) |
| URA3 | P03962 | 3gdl | GLN 215 | Transition state | BRENDA (*17*) |
| URA3 | P03962 | 3gdl | TYR 217 | Substrate binding | UniProt entry |
| URA3 | P03962 | 3gdl | ARG 235 | Substrate binding | UniProt entry |
| T7 RNAP | P00573 | 1qln | LYS 93 | Interface with DNA | Structural analysis |

| | | | | | |
|---|---|---|---|---|---|
| T7 RNAP | P00573 | 1qln | ALA 94 | Interface with DNA | Structural analysis |
| T7 RNAP | P00573 | 1qln | LYS 95 | Interface with DNA | Structural analysis |
| T7 RNAP | P00573 | 1qln | ARG 96 | Interface with DNA | Structural analysis |
| T7 RNAP | P00573 | 1qln | GLY 97 | Interface with DNA | Structural analysis |
| T7 RNAP | P00573 | 1qln | LYS 98 | Interface with DNA | Structural analysis |
| T7 RNAP | P00573 | 1qln | ARG 99 | Interface with DNA | Structural analysis |
| T7 RNAP | P00573 | 1qln | PRO 100 | Interface with DNA | Structural analysis |
| T7 RNAP | P00573 | 1qln | THR 101 | Interface with DNA | Structural analysis |
| T7 RNAP | P00573 | 1qln | GLN 104 | Interface with DNA | Structural analysis |
| T7 RNAP | P00573 | 1qln | ASN 131 | Interface with DNA | Structural analysis |
| T7 RNAP | P00573 | 1qln | THR 133 | Interface with DNA | Structural analysis |
| T7 RNAP | P00573 | 1qln | VAL 134 | Interface with DNA | Structural analysis |
| T7 RNAP | P00573 | 1qln | GLN 135 | Interface with DNA | Structural analysis |
| T7 RNAP | P00573 | 1qln | ALA 136 | Interface with DNA | Structural analysis |
| T7 RNAP | P00573 | 1qln | SER 139 | Interface with DNA | Structural analysis |
| T7 RNAP | P00573 | 1qln | ARG 143 | Interface with DNA | Structural analysis |
| T7 RNAP | P00573 | 1qln | ALA 200 | Interface with DNA | Structural analysis |
| T7 RNAP | P00573 | 1qln | TRP 201 | Interface with DNA | Structural analysis |
| T7 RNAP | P00573 | 1qln | LYS 206 | Interface with DNA | Structural analysis |
| T7 RNAP | P00573 | 1qln | ILE 210 | Interface with DNA | Structural analysis |
| T7 RNAP | P00573 | 1qln | HIS 211 | Interface with DNA | Structural analysis |
| T7 RNAP | P00573 | 1qln | ARG 215 | Interface with DNA | Structural analysis |

| | | | | | |
|---|---|---|---|---|---|
| T7 RNAP | P00573 | 1qln | ARG 231 | Interface with DNA | Structural analysis |
| T7 RNAP | P00573 | 1qln | ALA 234 | Interface with DNA | Structural analysis |
| T7 RNAP | P00573 | 1qln | GLY 235 | Interface with DNA | Structural analysis |
| T7 RNAP | P00573 | 1qln | VAL 237 | Interface with DNA | Structural analysis |
| T7 RNAP | P00573 | 1qln | GLY 238 | Interface with DNA | Structural analysis |
| T7 RNAP | P00573 | 1qln | ASP 240 | Interface with DNA | Structural analysis |
| T7 RNAP | P00573 | 1qln | SER 241 | Interface with DNA | Structural analysis |
| T7 RNAP | P00573 | 1qln | GLU 242 | Interface with DNA | Structural analysis |
| T7 RNAP | P00573 | 1qln | PHE 268 | Interface with DNA | Structural analysis |
| T7 RNAP | P00573 | 1qln | LEU 294 | Interface with DNA | Structural analysis |
| T7 RNAP | P00573 | 1qln | ALA 295 | Interface with DNA | Structural analysis |
| T7 RNAP | P00573 | 1qln | ARG 298 | Interface with DNA | Structural analysis |
| T7 RNAP | P00573 | 1qln | THR 299 | Interface with DNA | Structural analysis |
| T7 RNAP | P00573 | 1qln | HIS 300 | Interface with DNA | Structural analysis |
| T7 RNAP | P00573 | 1qln | ASN 419 | Interface with DNA | Structural analysis |
| T7 RNAP | P00573 | 1qln | ASP 421 | Interface with DNA | Structural analysis |
| T7 RNAP | P00573 | 1qln | TRP 422 | Interface with DNA | Structural analysis |
| T7 RNAP | P00573 | 1qln | ARG 423 | Interface with DNA | Structural analysis |
| T7 RNAP | P00573 | 1qln | ARG 425 | Interface with DNA | Structural analysis |
| T7 RNAP | P00573 | 1qln | TYR 427 | Interface with DNA | Structural analysis |
| T7 RNAP | P00573 | 1qln | GLN 435 | Interface with DNA | Structural analysis |
| T7 RNAP | P00573 | 1qln | GLY 436 | Interface with DNA | Structural analysis |

| | | | | | |
|---|---|---|---|---|---|
| T7 RNAP | P00573 | 1qln | ASN 437 | Interface with DNA | Structural analysis |
| T7 RNAP | P00573 | 1qln | LYS 441 | Interface with DNA | Structural analysis |
| T7 RNAP | P00573 | 1qln | ASP 537 | Active Site | UniProt entry |
| T7 RNAP | P00573 | 1qln | PRO 563 | Inactivated upon mutagenesis | UniProt entry |
| T7 RNAP | P00573 | 1qln | TYR 571 | Inactivated upon mutagenesis | UniProt entry |
| T7 RNAP | P00573 | 1qln | LYS 631 | Active site | UniProt entry |
| T7 RNAP | P00573 | 1qln | THR 636 | Inactivated upon mutagenesis | UniProt entry |
| T7 RNAP | P00573 | 1qln | TYR 639 | Inactivated upon mutagenesis | UniProt entry |
| T7 RNAP | P00573 | 1qln | GLY 640 | Interface with DNA | Structural analysis |
| T7 RNAP | P00573 | 1qln | SER 641 | Interface with DNA | Structural analysis |
| T7 RNAP | P00573 | 1qln | PHE 644 | Interface with DNA | Structural analysis |
| T7 RNAP | P00573 | 1qln | GLY 645 | Interface with DNA | Structural analysis |
| T7 RNAP | P00573 | 1qln | PHE 646 | Inactivated upon mutagenesis | UniProt entry |
| T7 RNAP | P00573 | 1qln | GLN 649 | Interface with DNA | Structural analysis |
| T7 RNAP | P00573 | 1qln | GLU 652 | Interface with DNA | Structural analysis |
| T7 RNAP | P00573 | 1qln | TYR 739 | Interface with DNA | Structural analysis |
| T7 RNAP | P00573 | 1qln | PRO 742 | Interface with DNA | Structural analysis |
| T7 RNAP | P00573 | 1qln | GLN 744 | Interface with DNA | Structural analysis |
| T7 RNAP | P00573 | 1qln | THR 745 | Interface with DNA | Structural analysis |
| T7 RNAP | P00573 | 1qln | ARG 746 | Interface with DNA | Structural analysis |
| T7 RNAP | P00573 | 1qln | LEU 747 | Interface with DNA | Structural analysis |
| T7 RNAP | P00573 | 1qln | ASN 748 | Interface with DNA | Structural analysis |

| | | | | | |
|---|---|---|---|---|---|
| T7 RNAP | P00573 | 1qln | MET 750 | Interface with DNA | Structural analysis |
| T7 RNAP | P00573 | 1qln | GLN 754 | Interface with DNA | Structural Analysis |
| T7 RNAP | P00573 | 1qln | PHE 755 | Interface with DNA | Structural analysis |
| T7 RNAP | P00573 | 1qln | ARG 756 | Interface with DNA | Structural analysis |
| T7 RNAP | P00573 | 1qln | LEU 757 | Interface with DNA | Structural analysis |
| T7 RNAP | P00573 | 1qln | GLN 758 | Interface with DNA | Structural analysis |
| T7 RNAP | P00573 | 1qln | PRO 759 | Interface with DNA | Structural analysis |
| T7 RNAP | P00573 | 1qln | THR 760 | Interface with DNA | Structural analysis |
| T7 RNAP | P00573 | 1qln | ILE 761 | Interface with DNA | Structural analysis |
| T7 RNAP | P00573 | 1qln | ASN 762 | Interface with DNA | Structural analysis |
| T7 RNAP | P00573 | 1qln | HIS 772 | Interface with DNA | Structural analysis |
| T7 RNAP | P00573 | 1qln | LYS 773 | Interface with DNA | Structural analysis |
| T7 RNAP | P00573 | 1qln | SER 776 | Interface with DNA | Structural analysis |
| T7 RNAP | P00573 | 1qln | GLY 777 | Interface with DNA | Structural analysis |
| T7 RNAP | P00573 | 1qln | PRO 780 | Interface with DNA | Structural analysis |
| T7 RNAP | P00573 | 1qln | ASN 781 | Interface with DNA | Structural analysis |
| T7 RNAP | P00573 | 1qln | HIS 784 | Interface with DNA | Structural analysis |
| T7 RNAP | P00573 | 1qln | ILE 810 | Interface with DNA | Structural analysis |
| T7 RNAP | P00573 | 1qln | HIS 811 | Interface with DNA | Structural analysis |
| T7 RNAP | P00573 | 1qln | ASP 812 | Active site | UniProt entry |

**Table S2.**

**NCBI amino acid BLAST+ results for synthetic homologs.** Sequences were submitted to the NCBI API using the default search parameters. The Expect- (E-) value is a statistical measure output from BLAST of the likelihood of finding the sequence in the database due to chance (9). Sequence identifiers: “psd95pdz3” = PDZ3, “ura3” = URA3, “t7rnapol” = T7 RNAP. Underscores indicate synthetic homolog number. Empty cells indicate no match found.

| Query sequence | Best match | E-value |
|---|---|---|
| PDZ3_0 | pdb\|6QJI\|A Chain A, Disks large homolog 4 [Homo sapiens]<br>>pdb\|6QJI\|B Chain B, Disks large homolog 4 [Homo sapiens]<br>>pdb\|6QJI\|C Chain C, Disks large homolog 4 [Homo sapiens]<br>>pdb\|6QJI\|D Chain D, Disks large homolog 4 [Homo sapiens]<br>>pdb\|6QJI\|E Chain E, Disks large homolog 4 [Homo sapiens]<br>>pdb\|6QJI\|F Chain F, Disks large homolog 4 [Homo sapiens]<br>>pdb\|6QJJ\|A Chain A, Disks large homolog 4 [Homo sapiens] | 3.0E-49 |
| PDZ3_1 | pdb\|6QJI\|A Chain A, Disks large homolog 4 [Homo sapiens]<br>>pdb\|6QJI\|B Chain B, Disks large homolog 4 [Homo sapiens]<br>>pdb\|6QJI\|C Chain C, Disks large homolog 4 [Homo sapiens]<br>>pdb\|6QJI\|D Chain D, Disks large homolog 4 [Homo sapiens]<br>>pdb\|6QJI\|E Chain E, Disks large homolog 4 [Homo sapiens]<br>>pdb\|6QJI\|F Chain F, Disks large homolog 4 [Homo sapiens]<br>>pdb\|6QJJ\|A Chain A, Disks large homolog 4 [Homo sapiens] | 2.5E-48 |
| PDZ3_2 | pdb\|6QJI\|A Chain A, Disks large homolog 4 [Homo sapiens]<br>>pdb\|6QJI\|B Chain B, Disks large homolog 4 [Homo sapiens]<br>>pdb\|6QJI\|C Chain C, Disks large homolog 4 [Homo sapiens]<br>>pdb\|6QJI\|D Chain D, Disks large homolog 4 [Homo sapiens]<br>>pdb\|6QJI\|E Chain E, Disks large homolog 4 [Homo sapiens]<br>>pdb\|6QJI\|F Chain F, Disks large homolog 4 [Homo sapiens]<br>>pdb\|6QJJ\|A Chain A, Disks large homolog 4 [Homo sapiens] | 6.9E-47 |
| PDZ3_3 | pdb\|6QJI\|A Chain A, Disks large homolog 4 [Homo sapiens]<br>>pdb\|6QJI\|B Chain B, Disks large homolog 4 [Homo sapiens]<br>>pdb\|6QJI\|C Chain C, Disks large homolog 4 [Homo sapiens]<br>>pdb\|6QJI\|D Chain D, Disks large homolog 4 [Homo sapiens]<br>>pdb\|6QJI\|E Chain E, Disks large homolog 4 [Homo sapiens]<br>>pdb\|6QJI\|F Chain F, Disks large homolog 4 [Homo sapiens]<br>>pdb\|6QJJ\|A Chain A, Disks large homolog 4 [Homo sapiens] | 5.6E-47 |
| PDZ3_4 | pdb\|6QJI\|A Chain A, Disks large homolog 4 [Homo sapiens]<br>>pdb\|6QJI\|B Chain B, Disks large homolog 4 [Homo sapiens]<br>>pdb\|6QJI\|C Chain C, Disks large homolog 4 [Homo sapiens]<br>>pdb\|6QJI\|D Chain D, Disks large homolog 4 [Homo sapiens]<br>>pdb\|6QJI\|E Chain E, Disks large homolog 4 [Homo sapiens]<br>>pdb\|6QJI\|F Chain F, Disks large homolog 4 [Homo sapiens]<br>>pdb\|6QJJ\|A Chain A, Disks large homolog 4 [Homo sapiens] | 4.6E-43 |
| PDZ3_5 | pdb\|6QJI\|A Chain A, Disks large homolog 4 [Homo sapiens]<br>>pdb\|6QJI\|B Chain B, Disks large homolog 4 [Homo sapiens] | 1.2E-44 |

| | | |
|---|---|---|
| | >pdb\|6QJI\|C Chain C, Disks large homolog 4 [Homo sapiens] >pdb\|6QJI\|D Chain D, Disks large homolog 4 [Homo sapiens] >pdb\|6QJI\|E Chain E, Disks large homolog 4 [Homo sapiens] >pdb\|6QJI\|F Chain F, Disks large homolog 4 [Homo sapiens] >pdb\|6QJJ\|A Chain A, Disks large homolog 4 [Homo sapiens] | |
| PDZ3_6 | pdb\|8AH4\|A Chain A, cDNA FLJ50577, highly similar to Discs large homolog 4 [Homo sapiens] >pdb\|8AH4\|B Chain B, cDNA FLJ50577, highly similar to Discs large homolog 4 [Homo sapiens] >pdb\|8AH4\|C Chain C, cDNA FLJ50577, highly similar to Discs large homolog 4 [Homo sapiens] >pdb\|8AH4\|D Chain D, cDNA FLJ50577, highly similar to Discs large homolog 4 [Homo sapiens] >pdb\|8AH4\|E Chain E, cDNA FLJ50577, highly similar to Discs large homolog 4 [Homo sapiens] >pdb\|8AH4\|F Chain F, cDNA FLJ50577, highly similar to Discs large homolog 4 [Homo sapiens] >pdb\|8AH5\|A Chain A, cDNA FLJ50577, highly similar to Discs large homolog 4 [Homo sapiens] >pdb\|8AH6\|A Chain A, cDNA FLJ50577, highly similar to Discs large homolog 4 [Homo sapiens] >pdb\|8AH6\|B Chain B, cDNA FLJ50577, highly similar to Discs large homolog 4 [Homo sapiens] >pdb\|8AH7\|A Chain A, cDNA FLJ50577, highly similar to Discs large homolog 4 [Homo sapiens] | 4.3E-44 |
| PDZ3_7 | pdb\|6QJI\|A Chain A, Disks large homolog 4 [Homo sapiens] >pdb\|6QJI\|B Chain B, Disks large homolog 4 [Homo sapiens] >pdb\|6QJI\|C Chain C, Disks large homolog 4 [Homo sapiens] >pdb\|6QJI\|D Chain D, Disks large homolog 4 [Homo sapiens] >pdb\|6QJI\|E Chain E, Disks large homolog 4 [Homo sapiens] >pdb\|6QJI\|F Chain F, Disks large homolog 4 [Homo sapiens] >pdb\|6QJJ\|A Chain A, Disks large homolog 4 [Homo sapiens] | 2.3E-43 |
| PDZ3_8 | pdb\|8AH4\|A Chain A, cDNA FLJ50577, highly similar to Discs large homolog 4 [Homo sapiens] >pdb\|8AH4\|B Chain B, cDNA FLJ50577, highly similar to Discs large homolog 4 [Homo sapiens] >pdb\|8AH4\|C Chain C, cDNA FLJ50577, highly similar to Discs large homolog 4 [Homo sapiens] >pdb\|8AH4\|D Chain D, cDNA FLJ50577, highly similar to Discs large homolog 4 [Homo sapiens] >pdb\|8AH4\|E Chain E, cDNA FLJ50577, highly similar to Discs large homolog 4 [Homo sapiens] >pdb\|8AH4\|F Chain F, cDNA FLJ50577, highly similar to Discs large homolog 4 [Homo sapiens] >pdb\|8AH5\|A Chain A, cDNA FLJ50577, highly similar to Discs large homolog 4 [Homo sapiens] >pdb\|8AH6\|A Chain A, cDNA FLJ50577, highly similar to Discs large homolog 4 [Homo sapiens] >pdb\|8AH6\|B Chain B, cDNA FLJ50577, highly similar to Discs large homolog 4 [Homo sapiens] >pdb\|8AH7\|A Chain A, cDNA FLJ50577, highly similar to Discs large homolog 4 [Homo sapiens] | 8.4E-45 |
| PDZ3_9 | pdb\|6QJI\|A Chain A, Disks large homolog 4 [Homo sapiens] >pdb\|6QJI\|B Chain B, Disks large homolog 4 [Homo sapiens] | 5.5E-43 |

| | | |
|---|---|---|
| | >pdb\|6QJI\|C Chain C, Disks large homolog 4 [Homo sapiens]<br>>pdb\|6QJI\|D Chain D, Disks large homolog 4 [Homo sapiens]<br>>pdb\|6QJI\|E Chain E, Disks large homolog 4 [Homo sapiens]<br>>pdb\|6QJI\|F Chain F, Disks large homolog 4 [Homo sapiens]<br>>pdb\|6QJJ\|A Chain A, Disks large homolog 4 [Homo sapiens] | |
| PDZ3_10 | pdb\|6QJD\|A Chain A, Disks large homolog 4 [Homo sapiens]<br>>pdb\|6QJD\|B Chain B, Disks large homolog 4 [Homo sapiens]<br>>pdb\|6QJD\|C Chain C, Disks large homolog 4 [Homo sapiens]<br>>pdb\|6QJD\|D Chain D, Disks large homolog 4 [Homo sapiens] | 4.7E-41 |
| PDZ3_11 | pdb\|6QJK\|A Chain A, Disks large homolog 4 [Homo sapiens] | 2.4E-35 |
| PDZ3_12 | pdb\|6QJK\|A Chain A, Disks large homolog 4 [Homo sapiens] | 2.4E-35 |
| PDZ3_13 | pdb\|6QJK\|A Chain A, Disks large homolog 4 [Homo sapiens] | 3.5E-36 |
| PDZ3_14 | pdb\|6QJK\|A Chain A, Disks large homolog 4 [Homo sapiens] | 9.5E-36 |
| PDZ3_15 | pdb\|6QJK\|A Chain A, Disks large homolog 4 [Homo sapiens] | 8.4E-35 |
| PDZ3_16 | pdb\|6QJK\|A Chain A, Disks large homolog 4 [Homo sapiens] | 8.4E-35 |
| PDZ3_17 | pdb\|6QJD\|A Chain A, Disks large homolog 4 [Homo sapiens]<br>>pdb\|6QJD\|B Chain B, Disks large homolog 4 [Homo sapiens]<br>>pdb\|6QJD\|C Chain C, Disks large homolog 4 [Homo sapiens]<br>>pdb\|6QJD\|D Chain D, Disks large homolog 4 [Homo sapiens] | 1.2E-38 |
| PDZ3_18 | pdb\|6QJI\|A Chain A, Disks large homolog 4 [Homo sapiens]<br>>pdb\|6QJI\|B Chain B, Disks large homolog 4 [Homo sapiens]<br>>pdb\|6QJI\|C Chain C, Disks large homolog 4 [Homo sapiens]<br>>pdb\|6QJI\|D Chain D, Disks large homolog 4 [Homo sapiens]<br>>pdb\|6QJI\|E Chain E, Disks large homolog 4 [Homo sapiens]<br>>pdb\|6QJI\|F Chain F, Disks large homolog 4 [Homo sapiens]<br>>pdb\|6QJJ\|A Chain A, Disks large homolog 4 [Homo sapiens] | 4.5E-39 |
| PDZ3_19 | pdb\|6QJD\|A Chain A, Disks large homolog 4 [Homo sapiens]<br>>pdb\|6QJD\|B Chain B, Disks large homolog 4 [Homo sapiens]<br>>pdb\|6QJD\|C Chain C, Disks large homolog 4 [Homo sapiens]<br>>pdb\|6QJD\|D Chain D, Disks large homolog 4 [Homo sapiens] | 7.0E-34 |
| PDZ3_20 | pdb\|6QJI\|A Chain A, Disks large homolog 4 [Homo sapiens]<br>>pdb\|6QJI\|B Chain B, Disks large homolog 4 [Homo sapiens]<br>>pdb\|6QJI\|C Chain C, Disks large homolog 4 [Homo sapiens]<br>>pdb\|6QJI\|D Chain D, Disks large homolog 4 [Homo sapiens]<br>>pdb\|6QJI\|E Chain E, Disks large homolog 4 [Homo sapiens]<br>>pdb\|6QJI\|F Chain F, Disks large homolog 4 [Homo sapiens]<br>>pdb\|6QJJ\|A Chain A, Disks large homolog 4 [Homo sapiens] | 4.5E-35 |
| PDZ3_21 | pdb\|2HE2\|A Chain A, Discs large homolog 2 [Homo sapiens]<br>>pdb\|2HE2\|B Chain B, Discs large homolog 2 [Homo sapiens] | 2.3E-34 |
| PDZ3_22 | pdb\|6QJK\|A Chain A, Disks large homolog 4 [Homo sapiens] | 3.7E-26 |
| PDZ3_23 | pdb\|6QJK\|A Chain A, Disks large homolog 4 [Homo sapiens] | 7.5E-27 |
| PDZ3_24 | pdb\|6QJK\|A Chain A, Disks large homolog 4 [Homo sapiens] | 4.9E-27 |
| PDZ3_25 | pdb\|6QJK\|A Chain A, Disks large homolog 4 [Homo sapiens] | 6.7E-23 |
| PDZ3_26 | gb\|KAH9383335.1\| hypothetical protein HPB48_024550 [Haemaphysalis longicornis] | 4.2E-26 |

| | | |
|---|---|---|
| PDZ3_27 | pdb\|6QJK\|A Chain A, Disks large homolog 4 [Homo sapiens] | 4.7E-22 |
| PDZ3_28 | pdb\|6QJI\|A Chain A, Disks large homolog 4 [Homo sapiens] >pdb\|6QJI\|B Chain B, Disks large homolog 4 [Homo sapiens] >pdb\|6QJI\|C Chain C, Disks large homolog 4 [Homo sapiens] >pdb\|6QJI\|D Chain D, Disks large homolog 4 [Homo sapiens] >pdb\|6QJI\|E Chain E, Disks large homolog 4 [Homo sapiens] >pdb\|6QJI\|F Chain F, Disks large homolog 4 [Homo sapiens] >pdb\|6QJJ\|A Chain A, Disks large homolog 4 [Homo sapiens] | 2.7E-28 |
| PDZ3_29 | gb\|KFV53693.1\| Disks large 4, partial [Gavia stellata] | 1.3E-29 |
| PDZ3_30 | pdb\|6QJK\|A Chain A, Disks large homolog 4 [Homo sapiens] | 1.1E-23 |
| PDZ3_31 | pdb\|6QJK\|A Chain A, Disks large homolog 4 [Homo sapiens] | 1.4E-19 |
| PDZ3_32 | pdb\|8AH4\|A Chain A, cDNA FLJ50577, highly similar to Discs large homolog 4 [Homo sapiens] >pdb\|8AH4\|B Chain B, cDNA FLJ50577, highly similar to Discs large homolog 4 [Homo sapiens] >pdb\|8AH4\|C Chain C, cDNA FLJ50577, highly similar to Discs large homolog 4 [Homo sapiens] >pdb\|8AH4\|D Chain D, cDNA FLJ50577, highly similar to Discs large homolog 4 [Homo sapiens] >pdb\|8AH4\|E Chain E, cDNA FLJ50577, highly similar to Discs large homolog 4 [Homo sapiens] >pdb\|8AH4\|F Chain F, cDNA FLJ50577, highly similar to Discs large homolog 4 [Homo sapiens] >pdb\|8AH5\|A Chain A, cDNA FLJ50577, highly similar to Discs large homolog 4 [Homo sapiens] >pdb\|8AH6\|A Chain A, cDNA FLJ50577, highly similar to Discs large homolog 4 [Homo sapiens] >pdb\|8AH6\|B Chain B, cDNA FLJ50577, highly similar to Discs large homolog 4 [Homo sapiens] >pdb\|8AH7\|A Chain A, cDNA FLJ50577, highly similar to Discs large homolog 4 [Homo sapiens] | 2.6E-29 |
| PDZ3_33 | gb\|NXB94462.1\| DLG4 protein [Vidua chalybeata] | 1.1E-29 |
| PDZ3_34 | pdb\|6QJK\|A Chain A, Disks large homolog 4 [Homo sapiens] | 2.6E-21 |
| PDZ3_35 | gb\|NXB73144.1\| DLG2 protein [Donacobius atricapilla] | 1.2E-26 |
| PDZ3_36 | pdb\|6QJK\|A Chain A, Disks large homolog 4 [Homo sapiens] | 6.0E-22 |
| PDZ3_37 | gb\|NXB94462.1\| DLG4 protein [Vidua chalybeata] | 3.5E-27 |
| PDZ3_38 | gb\|MEQ2182244.1\| Disks large 4 [Goodea atripinnis] | 6.6E-27 |
| PDZ3_39 | pdb\|8AH4\|A Chain A, cDNA FLJ50577, highly similar to Discs large homolog 4 [Homo sapiens] >pdb\|8AH4\|B Chain B, cDNA FLJ50577, highly similar to Discs large homolog 4 [Homo sapiens] >pdb\|8AH4\|C Chain C, cDNA FLJ50577, highly similar to Discs large homolog 4 [Homo sapiens] >pdb\|8AH4\|D Chain D, cDNA FLJ50577, highly similar to Discs large homolog 4 [Homo sapiens] >pdb\|8AH4\|E Chain E, cDNA FLJ50577, highly similar to Discs large homolog 4 [Homo sapiens] >pdb\|8AH4\|F Chain F, cDNA FLJ50577, highly similar to Discs large homolog 4 [Homo sapiens] >pdb\|8AH5\|A Chain A, cDNA FLJ50577, highly similar to Discs large homolog 4 [Homo sapiens] >pdb\|8AH6\|A Chain A, cDNA FLJ50577, highly similar to Discs large homolog 4 [Homo | 5.0E-25 |

<table>
<tr><td></td><td>sapiens] >pdb|8AH6|B Chain B, cDNA FLJ50577, highly similar to Discs large homolog 4 [Homo sapiens] >pdb|8AH7|A Chain A, cDNA FLJ50577, highly similar to Discs large homolog 4 [Homo sapiens]</td><td></td></tr>
<tr><td>PDZ3_40</td><td>gb|TRZ00889.1| hypothetical protein DNTS_002691, partial [Danionella cerebrum]</td><td>8.8E-21</td></tr>
<tr><td>PDZ3_41</td><td>gb|NXS13312.1| DLG3 protein [Neodrepanis coruscans] >gb|NXY25428.1| DLG3 protein [Atrichornis clamosus]</td><td>5.0E-22</td></tr>
<tr><td>PDZ3_42</td><td>gb|KAG8137043.1| hypothetical protein E2320_005588 [Naja naja]</td><td>3.0E-14</td></tr>
<tr><td>PDZ3_43</td><td>ref|XP_013187268.1| disks large 1 tumor suppressor protein isoform X4 [Amyelois transitella]</td><td>2.3E-16</td></tr>
<tr><td>PDZ3_44</td><td>ref|XP_030757886.1| disks large 1 tumor suppressor protein isoform X8 [Sitophilus oryzae]</td><td>2.6E-16</td></tr>
<tr><td>PDZ3_45</td><td>ref|XP_009047620.1| hypothetical protein LOTGIDRAFT_59096, partial [Lottia gigantea] >gb|ESP01730.1| hypothetical protein LOTGIDRAFT_59096, partial [Lottia gigantea]</td><td>5.5E-11</td></tr>
<tr><td>PDZ3_46</td><td>gb|KFV53693.1| Disks large 4, partial [Gavia stellata]</td><td>2.1E-13</td></tr>
<tr><td>PDZ3_47</td><td>ref|XP_044762455.1| disks large 1 tumor suppressor protein isoform X7 [Coccinella septempunctata]</td><td>2.4E-08</td></tr>
<tr><td>PDZ3_48</td><td>gb|NXM01890.1| DLG4 protein [Tyrannus savana] >gb|NXM24701.1| DLG4 protein [Oxyruncus cristatus]</td><td>3.7E-15</td></tr>
<tr><td>PDZ3_49</td><td>ref|XP_022242878.1| disks large homolog 1-like isoform X8 [Limulus polyphemus]</td><td>3.6E-12</td></tr>
<tr><td>PDZ3_50</td><td>ref|XP_065575668.1| disks large 1 tumor suppressor protein-like isoform X9 [Artemia franciscana]</td><td>7.3E-12</td></tr>
<tr><td>PDZ3_51</td><td>pdb|6QJK|A Chain A, Disks large homolog 4 [Homo sapiens]</td><td>7.0E-07</td></tr>
<tr><td>PDZ3_52</td><td>gb|KAL2301983.1| hypothetical protein Nmel_011385, partial [Mimus melanotis]</td><td>2.1E-12</td></tr>
<tr><td>PDZ3_53</td><td>ref|XP_033835444.1| tyrosine-protein phosphatase non-receptor type 13 [Periophthalmus magnuspinnatus]</td><td>1.1E-09</td></tr>
<tr><td>PDZ3_54</td><td>gb|KAJ8023385.1| Protein lin-7-like B [Holothuria leucospilota]</td><td>4.5E-12</td></tr>
<tr><td>PDZ3_55</td><td>emb|CAH1245870.1| DLG1 [Branchiostoma lanceolatum]</td><td>1.4E-02</td></tr>
<tr><td>PDZ3_56</td><td>ref|XP_033115095.1| multiple PDZ domain protein-like isoform X6 [Anneissia japonica]</td><td>7.5E-07</td></tr>
<tr><td>PDZ3_57</td><td>emb|CAF96151.1| unnamed protein product, partial [Tetraodon nigroviridis]</td><td>3.5E-11</td></tr>
<tr><td>PDZ3_58</td><td>ref|XP_034386756.1| Na(+)/H(+) exchange regulatory cofactor NHE-RF2 isoform X4 [Cyclopterus lumpus]</td><td>5.0E-06</td></tr>
<tr><td>PDZ3_59</td><td>ref|XP_065663661.1| tyrosine-protein phosphatase non-receptor type 13 isoform X3 [Hydra vulgaris]</td><td>1.5E-02</td></tr>
<tr><td>PDZ3_60</td><td>ref|XP_066993616.2| glutamate receptor-interacting protein 1 isoform X3 [Anabrus simplex]</td><td>8.2E-05</td></tr>
<tr><td>PDZ3_61</td><td>ref|XP_023311090.1| disks large 1 tumor suppressor protein isoform X7 [Anoplophora glabripennis]</td><td>1.1E-04</td></tr>
</table>

| | | |
|---|---|---|
| PDZ3_62 | emb\|CAD5116542.1\| DgyrCDS5421 [Dimorphilus gyrociliatus] | 5.7E-03 |
| PDZ3_63 | emb\|CAG5111368.1\| Oidioi.mRNA.OKI2018_I69.chr2.g5684.t1.cds [Oikopleura dioica] | 2.3E-01 |
| PDZ3_64 | gb\|KAG9259924.1\| multiple PDZ domain protein-like [Astyanax mexicanus] | 3.4E-04 |
| PDZ3_65 | emb\|CAH3146052.1\| unnamed protein product [Porites evermanni] | 4.4E-05 |
| PDZ3_66 | emb\|CAD2166676.1\| unnamed protein product [Meloidogyne enterolobii] | 1.3E-04 |
| PDZ3_67 | ref\|XP_048879408.1\| membrane-associated guanylate kinase, WW and PDZ domain-containing protein 3-like isoform X5 [Brienomyrus brachyistius] | 2.7E-12 |
| PDZ3_68 | ref\|XP_055618023.1\| disks large 1 tumor suppressor protein isoform X7 [Toxorhynchites rutilus septentrionalis] | 1.6E-06 |
| PDZ3_69 | NA | NA |
| PDZ3_70 | NA | NA |
| PDZ3_71 | NA | NA |
| PDZ3_72 | gb\|MBI4827283.1\| PAS domain S-box protein [Nitrospinota bacterium] | 7.3E-02 |
| PDZ3_73 | gb\|EJM27529.1\| hypothetical protein PMI24_03152 [Pseudomonas sp. GM25] | 3.9E+00 |
| PDZ3_74 | NA | NA |
| PDZ3_75 | ref\|WP_402137624.1\| nickel/cobalt transporter [Streptomyces sp. NPDC088727] >gb\|MFJ4898322.1\| nickel/cobalt transporter [Streptomyces sp. NPDC088727] | 4.8E+00 |
| PDZ3_76 | gb\|MCR4410444.1\| arabinose isomerase [Candidatus Saccharicenans sp.] >gb\|MDH7575952.1\| arabinose isomerase [Candidatus Saccharicenans sp.] | 6.1E+00 |
| PDZ3_77 | NA | NA |
| PDZ3_78 | gb\|MDQ7055620.1\| HTH domain-containing protein [Persephonella sp.] | 1.1E-04 |
| PDZ3_79 | NA | NA |
| PDZ3_80 | NA | NA |
| PDZ3_81 | NA | NA |
| PDZ3_82 | gb\|KAH8639486.1\| hypothetical protein IG631_07256 [Alternaria alternata] | 3.0E+00 |
| PDZ3_83 | NA | NA |
| PDZ3_84 | gb\|MEY2484583.1\| proton-dependent oligopeptide transporter, family [Verrucomicrobiota bacterium] | 3.9E+00 |
| PDZ3_85 | NA | NA |
| PDZ3_86 | tpg\|HTQ32706.1\| hydantoinase/oxoprolinase family protein [Stellaceae bacterium] | 9.0E+00 |
| PDZ3_87 | gb\|KFW89586.1\| Cytosolic carboxypeptidase 4, partial [Phalacrocorax carbo] | 1.9E-02 |

| | | |
|---|---|---|
| PDZ3_88 | NA | NA |
| PDZ3_89 | gb\|MCI6665562.1\| NUMOD4 motif-containing HNH endonuclease [Lachnospiraceae bacterium] | 4.5E-01 |
| PDZ3_90 | gb\|MDD3652693.1\| universal stress protein [Desulfotomaculaceae bacterium] | 8.2E+00 |
| PDZ3_91 | NA | NA |
| PDZ3_92 | NA | NA |
| PDZ3_93 | NA | NA |
| PDZ3_94 | NA | NA |
| PDZ3_95 | NA | NA |
| PDZ3_96 | NA | NA |
| PDZ3_97 | emb\|VDN07119.1\| unnamed protein product [Thelazia callipaeda] | 3.9E+00 |
| PDZ3_98 | NA | NA |
| PDZ3_99 | NA | NA |
| URA3_0 | emb\|CAD6622128.1\| XXYS1_4_G0003340.mRNA.1.CDS.1 [Saccharomyces cerevisiae] | 2.2E-136 |
| URA3_1 | emb\|CAD6622128.1\| XXYS1_4_G0003340.mRNA.1.CDS.1 [Saccharomyces cerevisiae] | 3.9E-133 |
| URA3_2 | dbj\|GHM90110.1\| orotidine 5'-phosphate decarboxylase [Saccharomyces cerevisiae] >emb\|CAI6461448.1\| CMF_HP2_G0014090.mRNA.1.CDS.1 [Saccharomyces cerevisiae] >emb\|CAI6470207.1\| CMF_HP1_G0014560.mRNA.1.CDS.1 [Saccharomyces cerevisiae] >emb\|CAI7265725.1\| CMF_collapsed_G0015770.mRNA.1.CDS.1 [Saccharomyces cerevisiae] | 8.1E-134 |
| URA3_3 | dbj\|GHM90110.1\| orotidine 5'-phosphate decarboxylase [Saccharomyces cerevisiae] >emb\|CAI6461448.1\| CMF_HP2_G0014090.mRNA.1.CDS.1 [Saccharomyces cerevisiae] >emb\|CAI6470207.1\| CMF_HP1_G0014560.mRNA.1.CDS.1 [Saccharomyces cerevisiae] >emb\|CAI7265725.1\| CMF_collapsed_G0015770.mRNA.1.CDS.1 [Saccharomyces cerevisiae] | 2.2E-129 |
| URA3_4 | dbj\|GHM90110.1\| orotidine 5'-phosphate decarboxylase [Saccharomyces cerevisiae] >emb\|CAI6461448.1\| CMF_HP2_G0014090.mRNA.1.CDS.1 [Saccharomyces cerevisiae] >emb\|CAI6470207.1\| CMF_HP1_G0014560.mRNA.1.CDS.1 [Saccharomyces cerevisiae] >emb\|CAI7265725.1\| CMF_collapsed_G0015770.mRNA.1.CDS.1 [Saccharomyces cerevisiae] | 6.8E-132 |
| URA3_5 | dbj\|GFP68107.1\| orotidine 5'-phosphate decarboxylase [Saccharomyces cerevisiae] >dbj\|GFP73067.1\| orotidine 5'-phosphate decarboxylase [Saccharomyces cerevisiae] | 2.8E-130 |

| | | |
|---|---|---|
| URA3_6 | ref\|XP_056081560.1\| orotidine-5'-phosphate decarboxylase [Saccharomyces mikatae IFO 1815] >emb\|CAI4038445.1\| hypothetical protein SMKI_05G0550 [Saccharomyces mikatae IFO 1815] | 8.2E-128 |
| URA3_7 | emb\|CAD6622128.1\| XXYS1_4_G0003340.mRNA.1.CDS.1 [Saccharomyces cerevisiae] | 3.0E-129 |
| URA3_8 | dbj\|GHM90110.1\| orotidine 5'-phosphate decarboxylase [Saccharomyces cerevisiae] >emb\|CAI6461448.1\| CMF_HP2_G0014090.mRNA.1.CDS.1 [Saccharomyces cerevisiae] >emb\|CAI6470207.1\| CMF_HP1_G0014560.mRNA.1.CDS.1 [Saccharomyces cerevisiae] >emb\|CAI7265725.1\| CMF_collapsed_G0015770.mRNA.1.CDS.1 [Saccharomyces cerevisiae] | 6.8E-122 |
| URA3_9 | dbj\|GHM90110.1\| orotidine 5'-phosphate decarboxylase [Saccharomyces cerevisiae] >emb\|CAI6461448.1\| CMF_HP2_G0014090.mRNA.1.CDS.1 [Saccharomyces cerevisiae] >emb\|CAI6470207.1\| CMF_HP1_G0014560.mRNA.1.CDS.1 [Saccharomyces cerevisiae] >emb\|CAI7265725.1\| CMF_collapsed_G0015770.mRNA.1.CDS.1 [Saccharomyces cerevisiae] | 1.2E-115 |
| URA3_10 | emb\|CAD6622128.1\| XXYS1_4_G0003340.mRNA.1.CDS.1 [Saccharomyces cerevisiae] | 6.5E-115 |
| URA3_11 | emb\|CAD6622128.1\| XXYS1_4_G0003340.mRNA.1.CDS.1 [Saccharomyces cerevisiae] | 2.1E-114 |
| URA3_12 | dbj\|GHM90110.1\| orotidine 5'-phosphate decarboxylase [Saccharomyces cerevisiae] >emb\|CAI6461448.1\| CMF_HP2_G0014090.mRNA.1.CDS.1 [Saccharomyces cerevisiae] >emb\|CAI6470207.1\| CMF_HP1_G0014560.mRNA.1.CDS.1 [Saccharomyces cerevisiae] >emb\|CAI7265725.1\| CMF_collapsed_G0015770.mRNA.1.CDS.1 [Saccharomyces cerevisiae] | 5.6E-118 |
| URA3_13 | emb\|CAH2352566.1\| orotidine 5'-phosphate decarboxylase [[Candida] railenensis] | 3.7E-111 |
| URA3_14 | emb\|CAH2352566.1\| orotidine 5'-phosphate decarboxylase [[Candida] railenensis] | 3.7E-105 |
| URA3_15 | ref\|XP_022465815.1\| orotidine-5'-phosphate decarboxylase [Huiozyma naganishii CBS 8797] >sp\|Q9Y726.1\| RecName: Full=Orotidine 5'-phosphate decarboxylase; AltName: Full=OMP decarboxylase; Short=OMPDCase; Short=OMPdecase; AltName: Full=Uridine 5'-monophosphate synthase; Short=UMP synthase [Maudiozyma exigua] >dbj\|BAA76736.1\| orotidine-5'-phosphate decarboxylase [Huiozyma naganishii] >emb\|CCK71570.1\| | 3.0E-102 |

<table>
<tr><td></td><td>hypothetical protein KNAG_0H01560 [Huiozyma naganishii CBS 8797]</td><td></td></tr>
<tr><td>URA3_16</td><td>ref|XP_064761013.1| orotidine-5'-phosphate decarboxylase [Scheffersomyces coipomensis] >gb|KAK6461656.1| orotidine-5'-phosphate decarboxylase [Scheffersomyces coipomensis]</td><td>6.1E-102</td></tr>
<tr><td>URA3_17</td><td>ref|XP_020061918.1| orotidine-5'-phosphate decarboxylase [Suhomyces tanzawaensis NRRL Y-17324] >gb|ODV76796.1| orotidine-5'-phosphate decarboxylase [Suhomyces tanzawaensis NRRL Y-17324]</td><td>2.5E-98</td></tr>
<tr><td>URA3_18</td><td>ref|XP_003677638.1| orotidine-5'-phosphate decarboxylase [Naumovozyma castellii] >emb|CCC71286.1| hypothetical protein NCAS_0G03990 [Naumovozyma castellii]</td><td>3.5E-98</td></tr>
<tr><td>URA3_19</td><td>emb|CAD6622128.1| XXYS1_4_G0003340.mRNA.1.CDS.1 [Saccharomyces cerevisiae]</td><td>4.1E-100</td></tr>
<tr><td>URA3_20</td><td>ref|XP_064761013.1| orotidine-5'-phosphate decarboxylase [Scheffersomyces coipomensis] >gb|KAK6461656.1| orotidine-5'-phosphate decarboxylase [Scheffersomyces coipomensis]</td><td>3.6E-96</td></tr>
<tr><td>URA3_21</td><td>ref|XP_056081560.1| orotidine-5'-phosphate decarboxylase [Saccharomyces mikatae IFO 1815] >emb|CAI4038445.1| hypothetical protein SMKI_05G0550 [Saccharomyces mikatae IFO 1815]</td><td>7.5E-100</td></tr>
<tr><td>URA3_22</td><td>ref|XP_056081560.1| orotidine-5'-phosphate decarboxylase [Saccharomyces mikatae IFO 1815] >emb|CAI4038445.1| hypothetical protein SMKI_05G0550 [Saccharomyces mikatae IFO 1815]</td><td>1.1E-106</td></tr>
<tr><td>URA3_23</td><td>gb|AJU49098.1| Ura3p [Saccharomyces cerevisiae YJM1383] >gb|AJU49606.1| Ura3p [Saccharomyces cerevisiae YJM1386] >gb|AJV33042.1| Ura3p [Saccharomyces cerevisiae YJM193] >gb|KZV11722.1| URA3 [Saccharomyces cerevisiae] >emb|CAI4403405.1| BCN_G0014650.mRNA.1.CDS.1 [Saccharomyces cerevisiae]</td><td>2.6E-97</td></tr>
<tr><td>URA3_24</td><td>gb|KAK1789115.1| hypothetical protein P4O66_015062 [Electrophorus voltai]</td><td>4.4E-95</td></tr>
<tr><td>URA3_25</td><td>gb|MDF2691546.1| pyrF [Gammaproteobacteria bacterium]</td><td>1.3E-94</td></tr>
<tr><td>URA3_26</td><td>emb|CAB1449281.1| unnamed protein product [Pleuronectes platessa]</td><td>3.3E-90</td></tr>
<tr><td>URA3_27</td><td>ref|XP_049340946.1| uridine 5'-monophosphate synthase [Astyanax mexicanus] >ref|XP_049340947.1| uridine 5'-monophosphate synthase [Astyanax mexicanus]</td><td>1.0E-85</td></tr>
<tr><td>URA3_28</td><td>emb|CAI4944458.1| CFC_HP_G0059700.mRNA.1.CDS.1 [Saccharomyces cerevisiae] >emb|CAI4951621.1| CFC_HP_G0064940.mRNA.1.CDS.1 [Saccharomyces cerevisiae] >emb|CAI6565978.1| CFC_HP_G0059700.mRNA.1.CDS.1 [Saccharomyces cerevisiae] >emb|CAI6614320.1| CFC_HP_G0064940.mRNA.1.CDS.1 [Saccharomyces cerevisiae]</td><td>6.2E-88</td></tr>
</table>

| | | |
|---|---|---|
| | >emb\|CAI7253598.1\| CFC_collapsed_G0014350.mRNA.1.CDS.1 [Saccharomyces cerevisiae] | |
| URA3_29 | ref\|XP_045535659.1\| uridine 5'-monophosphate synthase [Papilio machaon] | 4.6E-90 |
| URA3_30 | emb\|SCW02670.1\| LAFE_0F11738g1_1 [Lachancea fermentati] | 1.1E-82 |
| URA3_31 | ref\|XP_029320804.1\| uncharacterized protein C5L36_0B05730 [Pichia kudriavzevii] >sp\|Q6IUR4.1\| RecName: Full=Orotidine 5'-phosphate decarboxylase; AltName: Full=OMP decarboxylase; Short=OMPDCase; Short=OMPdecase; AltName: Full=Uridine 5'-monophosphate synthase; Short=UMP synthase [Pichia kudriavzevii] >gb\|QSG73538.1\| Ura3 [Cloning vector pWSPK-Cas9] >gb\|AAT39474.1\| orotidine-5'-monophosphate decarboxylase [Pichia kudriavzevii] >gb\|AWU75327.1\| hypothetical protein C5L36_0B05730 [Pichia kudriavzevii] >gb\|KGK39727.1\| hypothetical protein JL09_g1026 [Pichia kudriavzevii] >gb\|OUT21549.1\| orotidine 5'-phosphate decarboxylase [Pichia kudriavzevii] | 2.9E-73 |
| URA3_32 | emb\|SCW02670.1\| LAFE_0F11738g1_1 [Lachancea fermentati] | 1.2E-82 |
| URA3_33 | emb\|SCU98197.1\| LAFA_0G16248g1_1 [Lachancea sp. 'fantastica'] | 2.7E-71 |
| URA3_34 | emb\|SCU99172.1\| LADA_0H18008g1_1 [Lachancea dasiensis] | 4.3E-78 |
| URA3_35 | emb\|SCU99172.1\| LADA_0H18008g1_1 [Lachancea dasiensis] | 1.9E-82 |
| URA3_36 | sp\|Q12604.1\| RecName: Full=Orotidine 5'-phosphate decarboxylase; AltName: Full=OMP decarboxylase; Short=OMPDCase; Short=OMPdecase; AltName: Full=Uridine 5'-monophosphate synthase; Short=UMP synthase [Magnusiomyces magnusii] >emb\|CAA65135.1\| orotidine 5' phosphate decarboxylase [Magnusiomyces magnusii] | 1.6E-81 |
| URA3_37 | emb\|CAH0719549.1\| unnamed protein product, partial [Brenthis ino] | 3.8E-65 |
| URA3_38 | sp\|Q9HFX0.1\| RecName: Full=Orotidine 5'-phosphate decarboxylase; AltName: Full=OMP decarboxylase; Short=OMPDCase; Short=OMPdecase; AltName: Full=Uridine 5'-monophosphate synthase; Short=UMP synthase [Zygosaccharomyces bailii] >gb\|AAG17694.1\| orotidine-5'-phosphate decarboxylase [Zygosaccharomyces bailii] >emb\|CDF90219.1\| ZYBA0S06-03488g1_1 [Zygosaccharomyces bailii CLIB 213] >emb\|CDH11557.1\| Orotidine 5'-phosphate decarboxylase [Zygosaccharomyces bailii ISA1307] | 1.1E-77 |
| URA3_39 | gb\|PIY95553.1\| MAG: orotidine-5'-phosphate decarboxylase [Candidatus Kerfeldbacteria bacterium CG_4_10_14_0_8_um_filter_42_10] | 8.4E-62 |
| URA3_40 | gb\|OWK74209.1\| orotidine 5'-phosphate decarboxylase [Flavobacteriaceae bacterium JJC] | 1.3E-61 |

| | | |
|---|---|---|
| URA3_41 | gb\|MFH1426699.1\| orotidine-5'-phosphate decarboxylase [Candidatus Kerfeldbacteria bacterium] | 1.1E-59 |
| URA3_42 | ref\|XP_051619975.1\| URA3 [Candida jiufengensis] >gb\|KAI5953233.1\| URA3 [Candida jiufengensis] | 2.3E-66 |
| URA3_43 | ref\|XP_019017989.1\| hypothetical protein PICMEDRAFT_72906 [Pichia membranifaciens NRRL Y-2026] >gb\|ODQ46876.1\| hypothetical protein PICMEDRAFT_72906 [Pichia membranifaciens NRRL Y-2026] | 4.8E-60 |
| URA3_44 | gb\|XBW37801.1\| hypothetical protein QEN19_003378 [Hanseniaspora menglaensis] | 2.2E-61 |
| URA3_45 | gb\|OGT60575.1\| MAG: orotidine 5'-phosphate decarboxylase [Gammaproteobacteria bacterium RIFCSPHIGHO2_12_FULL_45_12] | 6.6E-58 |
| URA3_46 | ref\|XP_066795822.1\| Orotidine 5'-phosphate decarboxylase domain-containing protein [Lipomyces doorenjongii] >gb\|KAK9492897.1\| Orotidine 5'-phosphate decarboxylase domain-containing protein [Lipomyces doorenjongii] | 5.8E-37 |
| URA3_47 | dbj\|BAC20169.1\| orotidine-5'-phosphate decarboxylase [Millerozyma farinosa] | 2.1E-45 |
| URA3_48 | sp\|P78724.1\| RecName: Full=Orotidine 5'-phosphate decarboxylase; AltName: Full=OMP decarboxylase; Short=OMPDCase; Short=OMPdecase; AltName: Full=Uridine 5'-monophosphate synthase; Short=UMP synthase [Wickerhamomyces anomalus NRRL Y-366] >emb\|CAA70421.1\| orotidine-5'-phosphate decarboxylase [Wickerhamomyces anomalus] | 1.6E-45 |
| URA3_49 | gb\|KAF7278683.1\| hypothetical protein GWI33_008131 [Rhynchophorus ferrugineus] | 5.0E-15 |
| URA3_50 | ref\|XP_056081560.1\| orotidine-5'-phosphate decarboxylase [Saccharomyces mikatae IFO 1815] >emb\|CAI4038445.1\| hypothetical protein SMKI_05G0550 [Saccharomyces mikatae IFO 1815] | 6.9E-29 |
| URA3_51 | dbj\|BDT06277.1\| orotidine-5'-phosphate decarboxylase Ura3 [Expression vector YHp22601] >dbj\|BDT26484.1\| orotidine-5'-phosphate decarboxylase Ura3 [Expression vector YHp26352] | 1.3E-21 |
| URA3_52 | ref\|XP_022674242.1\| orotidine-5'-phosphate decarboxylase [Kluyveromyces marxianus DMKU3-1042] >sp\|P41769.1\| RecName: Full=Orotidine 5'-phosphate decarboxylase; AltName: Full=OMP decarboxylase; Short=OMPDCase; Short=OMPdecase; AltName: Full=Uridine 5'-monophosphate synthase; Short=UMP synthase [Kluyveromyces marxianus] >gb\|ABB69700.1\| orotidine-5'-phosphate decarboxylase [Kluyveromyces marxianus] >gb\|AOP17589.1\| orotidine-5'-phosphate decarboxylase [Kluyveromyces marxianus] >gb\|QGN13471.1\| Orotidine 5-phosphate decarboxylase [Kluyveromyces marxianus] >emb\|CAA79928.1\| URA3 [Kluyveromyces marxianus] | 1.6E-22 |

| | | |
|---|---|---|
| | >dbj\|BAO38353.1\| orotidine 5'-phosphate decarboxylase [Kluyveromyces marxianus DMKU3-1042] | |
| URA3_53 | ref\|XP_068758460.1\| uridine 5'-monophosphate synthase-like [Montipora capricornis] | 2.6E-21 |
| URA3_54 | emb\|CAI4944458.1\| CFC_HP_G0059700.mRNA.1.CDS.1 [Saccharomyces cerevisiae] >emb\|CAI4951621.1\| CFC_HP_G0064940.mRNA.1.CDS.1 [Saccharomyces cerevisiae] >emb\|CAI6565978.1\| CFC_HP_G0059700.mRNA.1.CDS.1 [Saccharomyces cerevisiae] >emb\|CAI6614320.1\| CFC_HP_G0064940.mRNA.1.CDS.1 [Saccharomyces cerevisiae] >emb\|CAI7253598.1\| CFC_collapsed_G0014350.mRNA.1.CDS.1 [Saccharomyces cerevisiae] | 1.8E-20 |
| URA3_55 | gb\|EJS44107.1\| ura3p [Saccharomyces arboricola H-6] | 6.9E-19 |
| URA3_56 | gb\|KAG0679187.1\| orotidine 5'-phosphate decarboxylase [Kluyveromyces marxianus] >gb\|KAG0685086.1\| orotidine 5'-phosphate decarboxylase [Kluyveromyces marxianus] | 4.5E-21 |
| URA3_57 | emb\|CDO95120.1\| unnamed protein product [Kluyveromyces dobzhanskii CBS 2104] | 2.9E-19 |
| URA3_58 | emb\|SCV04999.1\| LANO_0G16270g1_1 [Lachancea nothofagi CBS 11611] | 5.5E-15 |
| URA3_59 | ref\|XP_056081560.1\| orotidine-5'-phosphate decarboxylase [Saccharomyces mikatae IFO 1815] >emb\|CAI4038445.1\| hypothetical protein SMKI_05G0550 [Saccharomyces mikatae IFO 1815] | 6.5E-15 |
| URA3_60 | ref\|XP_003677638.1\| orotidine-5'-phosphate decarboxylase [Naumovozyma castellii] >emb\|CCC71286.1\| hypothetical protein NCAS_0G03990 [Naumovozyma castellii] | 3.4E-22 |
| URA3_61 | gb\|AGE89246.1\| orotidine-5'-phosphate decarboxylase [Schwanniomyces occidentalis] >dbj\|BAV32324.1\| orotidine-5'-phosphate decarboxylase [Schwanniomyces occidentalis var. occidentalis] | 1.0E-14 |
| URA3_62 | emb\|CAI5072685.1\| CRE_HP_G0117940.mRNA.1.CDS.1 [Saccharomyces cerevisiae] >emb\|CAI5157952.1\| CRE_HP_G0145160.mRNA.1.CDS.1 [Saccharomyces cerevisiae] >emb\|CAI6964968.1\| CRE_HP_G0117940.mRNA.1.CDS.1 [Saccharomyces cerevisiae] >emb\|CAI6993079.1\| CRE_HP_G0145160.mRNA.1.CDS.1 [Saccharomyces cerevisiae] | 1.1E-14 |
| URA3_63 | gb\|MBN2461395.1\| orotate phosphoribosyltransferase [Candidatus Cloacimonadota bacterium] | 4.2E-01 |
| URA3_64 | gb\|KAG0655378.1\| orotidine 5'-phosphate decarboxylase [Maudiozyma exigua] | 2.8E-18 |
| URA3_65 | gb\|WJX87251.1\| Uridine 5'-monophosphate synthase [Trifolium repens] | 2.4E-02 |
| URA3_66 | ref\|XP_033765792.1\| orotidine-5'-phosphate decarboxylase [Saccharomyces paradoxus] >gb\|QHS72760.1\| Ura3 [Saccharomyces paradoxus] | 1.2E-19 |

| | | |
|---|---|---|
| URA3_67 | gb\|QEU61253.1\| Ura3 [Kluyveromyces lactis] | 1.3E-15 |
| URA3_68 | ref\|XP_056087197.1\| orotidine-5'-phosphate decarboxylase [Saccharomyces kudriavzevii IFO 1802] >gb\|EJT41855.1\| URA3-like protein [Saccharomyces kudriavzevii IFO 1802] >emb\|CAI4059903.1\| hypothetical protein SKDI_05G0470 [Saccharomyces kudriavzevii IFO 1802] | 1.4E-19 |
| URA3_69 | ref\|XP_056087197.1\| orotidine-5'-phosphate decarboxylase [Saccharomyces kudriavzevii IFO 1802] >gb\|EJT41855.1\| URA3-like protein [Saccharomyces kudriavzevii IFO 1802] >emb\|CAI4059903.1\| hypothetical protein SKDI_05G0470 [Saccharomyces kudriavzevii IFO 1802] | 6.9E-23 |
| URA3_70 | emb\|SCU99172.1\| LADA_0H18008g1_1 [Lachancea dasiensis] | 1.2E-19 |
| URA3_71 | gb\|KAG0655378.1\| orotidine 5'-phosphate decarboxylase [Maudiozyma exigua] | 5.3E-19 |
| URA3_72 | emb\|SCV04999.1\| LANO_0G16270g1_1 [Lachancea nothofagi CBS 11611] | 1.9E-18 |
| URA3_73 | ref\|XP_033765792.1\| orotidine-5'-phosphate decarboxylase [Saccharomyces paradoxus] >gb\|QHS72760.1\| Ura3 [Saccharomyces paradoxus] | 1.4E-20 |
| URA3_74 | ref\|XP_018222653.1\| orotidine-5'-phosphate decarboxylase [Saccharomyces eubayanus] >gb\|QID84527.1\| orotidine 5'-phosphate decarboxylase [Saccharomyces pastorianus] >gb\|KOG99935.1\| URA3-like protein [Saccharomyces eubayanus] >emb\|CAI1945211.1\| hypothetical protein SEUBUCD650_0E00740 [Saccharomyces eubayanus] >emb\|CAI1974505.1\| hypothetical protein SEUBUCD646_0E00700 [Saccharomyces eubayanus] >dbj\|BAI82233.1\| orotidine-5'-phosphate decarboxylase [Saccharomyces pastorianus] | 1.0E-16 |
| URA3_75 | gb\|AJU49098.1\| Ura3p [Saccharomyces cerevisiae YJM1383] >gb\|AJU49606.1\| Ura3p [Saccharomyces cerevisiae YJM1386] >gb\|AJV33042.1\| Ura3p [Saccharomyces cerevisiae YJM193] >gb\|KZV11722.1\| URA3 [Saccharomyces cerevisiae] >emb\|CAI4403405.1\| BCN_G0014650.mRNA.1.CDS.1 [Saccharomyces cerevisiae] | 1.4E-15 |
| URA3_76 | NA | NA |
| URA3_77 | gb\|MCF8531851.1\| methylmalonyl-CoA carboxyltransferase [Reyranella sp.] | 6.1E-01 |
| URA3_78 | NA | NA |
| URA3_79 | emb\|CDO95120.1\| unnamed protein product [Kluyveromyces dobzhanskii CBS 2104] | 3.9E-25 |
| URA3_80 | NA | NA |
| URA3_81 | emb\|SCV04999.1\| LANO_0G16270g1_1 [Lachancea nothofagi CBS 11611] | 1.3E-14 |

| | | |
|---|---|---|
| URA3_82 | emb\|CAI7112224.1\| BAF_collapsed_G0014430.mRNA.1.CDS.1 [Saccharomyces cerevisiae] | 4.0E-17 |
| URA3_83 | NA | NA |
| URA3_84 | NA | NA |
| URA3_85 | gb\|TVS08297.1\| MAG: hypothetical protein EA417_23040 [Gammaproteobacteria bacterium] | 2.6E+00 |
| URA3_86 | NA | NA |
| URA3_87 | gb\|AJU50614.1\| Ura3p [Saccharomyces cerevisiae YJM1399] >emb\|CAI4403622.1\| CPI_1c_G0014250.mRNA.1.CDS.1 [Saccharomyces cerevisiae] >emb\|CAI4414414.1\| BBM_1a_G0014170.mRNA.1.CDS.1 [Saccharomyces cerevisiae] >emb\|CAI4416540.1\| ADE_G0014090.mRNA.1.CDS.1 [Saccharomyces cerevisiae] >emb\|CAI4418990.1\| CDN_1a_G0014280.mRNA.1.CDS.1 [Saccharomyces cerevisiae] | 1.3E-11 |
| URA3_88 | NA | NA |
| URA3_89 | gb\|KAK3717004.1\| orotidine 5'-phosphate decarboxylase [Vermiconidia calcicola] | 1.8E-01 |
| URA3_90 | gb\|RLD49595.1\| MAG: orotidine 5'-phosphate decarboxylase [Bacteroidota bacterium] | 3.0E+00 |
| URA3_91 | gb\|OGT60670.1\| MAG: orotidine 5'-phosphate decarboxylase [Gammaproteobacteria bacterium RIFCSPHIGHO2_12_FULL_43_28] | 1.9E-02 |
| URA3_92 | emb\|CDO95120.1\| unnamed protein product [Kluyveromyces dobzhanskii CBS 2104] | 1.6E-03 |
| URA3_93 | ref\|WP_149245606.1\| orotidine-5'-phosphate decarboxylase [Chryseobacterium sp. SN22] >gb\|KAA0130513.1\| orotate phosphoribosyltransferase [Chryseobacterium sp. SN22] | 5.3E+00 |
| URA3_94 | gb\|MCF6367177.1\| orotidine-5'-phosphate decarboxylase [Bacteroidales bacterium] | 6.0E+00 |
| URA3_95 | NA | NA |
| URA3_96 | gb\|KAK3171566.1\| hypothetical protein OEA41_003650 [Lepraria neglecta] | 5.6E-02 |
| URA3_97 | NA | NA |
| URA3_98 | NA | NA |
| URA3_99 | ref\|XP_031549189.1\| UHRF1-binding protein 1-like isoform X2 [Actinia tenebrosa] | 4.9E-02 |
| T7RNAP_0 | pdb\|1ARO\|P Chain P, T7 RNA POLYMERASE [Escherichia phage T7] | 0 |
| T7RNAP_1 | ref\|WP_260790296.1\| DNA-directed RNA polymerase [Pseudomonas aeruginosa] >gb\|UXA41723.1\| hypothetical protein MMG97_16465 [Pseudomonas aeruginosa] | 0 |
| T7RNAP_2 | pdb\|1ARO\|P Chain P, T7 RNA POLYMERASE [Escherichia phage T7] | 0 |
| T7RNAP_3 | gb\|WPK33069.1\| DNA-directed RNA polymerase [Escherichia phage AV103] | 0 |

| | | |
|---|---|---|
| T7RNAP_4 | pdb\|1ARO\|P Chain P, T7 RNA POLYMERASE [Escherichia phage T7] | 0 |
| T7RNAP_5 | ref\|WP_260790296.1\| DNA-directed RNA polymerase [Pseudomonas aeruginosa] >gb\|UXA41723.1\| hypothetical protein MMG97_16465 [Pseudomonas aeruginosa] | 0 |
| T7RNAP_6 | gb\|WPK33069.1\| DNA-directed RNA polymerase [Escherichia phage AV103] | 0 |
| T7RNAP_7 | pdb\|1ARO\|P Chain P, T7 RNA POLYMERASE [Escherichia phage T7] | 0 |
| T7RNAP_8 | pdb\|1ARO\|P Chain P, T7 RNA POLYMERASE [Escherichia phage T7] | 0 |
| T7RNAP_9 | pdb\|1ARO\|P Chain P, T7 RNA POLYMERASE [Escherichia phage T7] | 0 |
| T7RNAP_10 | pdb\|1ARO\|P Chain P, T7 RNA POLYMERASE [Escherichia phage T7] | 0 |
| T7RNAP_11 | gb\|AAA32569.1\| RNA polymerase [Escherichia phage T7] >gb\|ALY06011.1\| T7 RNA polymerase [Cloning vector pAKT7] >gb\|ALY06013.1\| T7 RNA polymerase [Cloning vector pYAT7] | 0 |
| T7RNAP_12 | gb\|XLQ29395.1\| RNA polymerase [Escherichia phage KKE5P] | 0 |
| T7RNAP_13 | gb\|QXV80637.1\| RNA polymerase [Escherichia phage JacobBurckhardt] | 0 |
| T7RNAP_14 | pdb\|1ARO\|P Chain P, T7 RNA POLYMERASE [Escherichia phage T7] | 0 |
| T7RNAP_15 | ref\|YP_009804754.1\| RNA polymerase [Salmonella phage 3A_8767] >gb\|AXC37072.1\| DNA-directed RNA polymerase [Salmonella phage 3A_8767] | 0 |
| T7RNAP_16 | pdb\|1ARO\|P Chain P, T7 RNA POLYMERASE [Escherichia phage T7] | 0 |
| T7RNAP_17 | gb\|AAA32569.1\| RNA polymerase [Escherichia phage T7] >gb\|ALY06011.1\| T7 RNA polymerase [Cloning vector pAKT7] >gb\|ALY06013.1\| T7 RNA polymerase [Cloning vector pYAT7] | 0 |
| T7RNAP_18 | pdb\|1ARO\|P Chain P, T7 RNA POLYMERASE [Escherichia phage T7] | 0 |
| T7RNAP_19 | gb\|WPK33069.1\| DNA-directed RNA polymerase [Escherichia phage AV103] | 0 |
| T7RNAP_20 | pdb\|1ARO\|P Chain P, T7 RNA POLYMERASE [Escherichia phage T7] | 0 |
| T7RNAP_21 | gb\|XLQ29395.1\| RNA polymerase [Escherichia phage KKE5P] | 0 |
| T7RNAP_22 | ref\|WP_260790296.1\| DNA-directed RNA polymerase [Pseudomonas aeruginosa] >gb\|UXA41723.1\| hypothetical protein MMG97_16465 [Pseudomonas aeruginosa] | 0 |
| T7RNAP_23 | gb\|WPK33069.1\| DNA-directed RNA polymerase [Escherichia phage AV103] | 0 |
| T7RNAP_24 | pdb\|1ARO\|P Chain P, T7 RNA POLYMERASE [Escherichia phage T7] | 0 |

| | | |
|---|---|---|
| T7RNAP_25 | gb\|WPK33069.1\| DNA-directed RNA polymerase [Escherichia phage AV103] | 0 |
| T7RNAP_26 | ref\|WP_260790296.1\| DNA-directed RNA polymerase [Pseudomonas aeruginosa] >gb\|UXA41723.1\| hypothetical protein MMG97_16465 [Pseudomonas aeruginosa] | 0 |
| T7RNAP_27 | pdb\|1ARO\|P Chain P, T7 RNA POLYMERASE [Escherichia phage T7] | 0 |
| T7RNAP_28 | gb\|WPK33069.1\| DNA-directed RNA polymerase [Escherichia phage AV103] | 0 |
| T7RNAP_29 | gb\|AAA32569.1\| RNA polymerase [Escherichia phage T7] >gb\|ALY06011.1\| T7 RNA polymerase [Cloning vector pAKT7] >gb\|ALY06013.1\| T7 RNA polymerase [Cloning vector pYAT7] | 0 |
| T7RNAP_30 | gb\|WEU68248.1\| DNA-directed RNA polymerase [Escherichia phage vB_Ec_Tarrare] | 1.9E-132 |
| T7RNAP_31 | pdb\|1ARO\|P Chain P, T7 RNA POLYMERASE [Escherichia phage T7] | 5.8E-139 |
| T7RNAP_32 | gb\|QWY13933.1\| T3/T7-like RNA polymerase [Escherichia phage T7] >gb\|QWY13983.1\| T3/T7-like RNA polymerase [Escherichia phage T7] | 1.3E-147 |
| T7RNAP_33 | gb\|WBF69716.1\| DNA-directed RNA polymerase [Shigella phage SFP21B] | 9.1E-77 |
| T7RNAP_34 | gb\|QLF85705.1\| DNA-directed RNA polymerase [Serratia phage vB_SmaP-UFV01] | 4.3E-123 |
| T7RNAP_35 | gb\|AZV02339.2\| DNA-directed RNA polymerase [Pectobacterium phage Q19] >gb\|WOL25676.1\| hypothetical protein [Pectobacterium phage PcaP1EGY] | 3.9E-84 |
| T7RNAP_36 | gb\|QOV06378.1\| DNA-directed RNA polymerase [Escherichia phage JB01] | 4.7E-141 |
| T7RNAP_37 | emb\|CAA24333.1\| unnamed protein product [Escherichia phage T7] | 3.1E-129 |
| T7RNAP_38 | gb\|WEU68248.1\| DNA-directed RNA polymerase [Escherichia phage vB_Ec_Tarrare] | 1.2E-123 |
| T7RNAP_39 | gb\|WPJ67457.1\| RNA polymerase [Escherichia phage Carena] | 7.6E-81 |
| T7RNAP_40 | tpg\|HCU2121986.1\| DNA-dependent RNA polymerase [Klebsiella pneumoniae] | 5.8E-90 |
| T7RNAP_41 | gb\|UJQ71098.1\| RNA polymerase [Escherichia phage T7] | 4.3E-72 |
| T7RNAP_42 | gb\|QWY13933.1\| T3/T7-like RNA polymerase [Escherichia phage T7] >gb\|QWY13983.1\| T3/T7-like RNA polymerase [Escherichia phage T7] | 1.9E-136 |
| T7RNAP_43 | gb\|KAL2513886.1\| Leucine-rich repeat receptor-like serine/threonine-protein kinase BAM3 [Forsythia ovata] | 1.6E-134 |
| T7RNAP_44 | pdb\|1ARO\|P Chain P, T7 RNA POLYMERASE [Escherichia phage T7] | 6.3E-117 |

| | | |
|---|---|---|
| T7RNAP_45 | gb\|AIS22793.1\| UmuD-T7 RNA polymerase(1-514) [synthetic vector pCOLA-AraC-pBAD-T7_RNAP(515-884)-UmuD-T7RNAP(1-514)] | 1.8E-57 |
| T7RNAP_46 | gb\|AIS22793.1\| UmuD-T7 RNA polymerase(1-514) [synthetic vector pCOLA-AraC-pBAD-T7_RNAP(515-884)-UmuD-T7RNAP(1-514)] | 9.4E-56 |
| T7RNAP_47 | gb\|MDA2621644.1\| hypothetical protein [Bacillus cereus] | 1.6E-49 |
| T7RNAP_48 | gb\|AIS22794.1\| T7 RNA polymerase (515-884) [synthetic vector pCOLA-AraC-pBAD-T7_RNAP(515-884)-UmuD-T7RNAP(1-514)] | 1.7E-48 |
| T7RNAP_49 | gb\|QHJ79918.1\| MAG: hypothetical protein [Caudoviricetes sp.] | 9.3E-52 |
| T7RNAP_50 | gb\|WCS66925.1\| DNA-directed RNA polymerase [Serratia phage HMGUsm2] | 1.7E-43 |
| T7RNAP_51 | NA | NA |
| T7RNAP_52 | gb\|MDA2621644.1\| hypothetical protein [Bacillus cereus] | 3.6E-28 |
| T7RNAP_53 | ref\|YP_009187276.1\| N4-like RNA polymerase [Yersinia phage vB_YenP_AP10] >gb\|ALK86936.1\| RNA polymerase [Yersinia phage vB_YenP_AP10] | 3.5E-53 |
| T7RNAP_54 | gb\|UTQ80063.1\| RNA polymerase [Erwinia phage Stepyanka] | 7.6E-05 |
| T7RNAP_55 | gb\|QZB84819.1\| RNA polymerase [Escherichia phage T7] | 6.3E-57 |
| T7RNAP_56 | gb\|WOG34502.1\| T7 RNA polymerase, partial [synthetic construct] >gb\|WOG34505.1\| T7 RNA polymerase, partial [synthetic construct] | 1.2E-47 |
| T7RNAP_57 | NA | NA |
| T7RNAP_58 | NA | NA |
| T7RNAP_59 | NA | NA |
| T7RNAP_60 | NA | NA |
| T7RNAP_61 | NA | NA |
| T7RNAP_62 | NA | NA |
| T7RNAP_63 | NA | NA |
| T7RNAP_64 | NA | NA |
| T7RNAP_65 | NA | NA |
| T7RNAP_66 | NA | NA |
| T7RNAP_67 | NA | NA |
| T7RNAP_68 | NA | NA |
| T7RNAP_69 | NA | NA |
| T7RNAP_70 | NA | NA |
| T7RNAP_71 | NA | NA |
| T7RNAP_72 | NA | NA |
| T7RNAP_73 | NA | NA |
| T7RNAP_74 | NA | NA |
| T7RNAP_75 | NA | NA |

| | | |
|---|---|---|
| T7RNAP_76 | NA | NA |
| T7RNAP_77 | NA | NA |
| T7RNAP_78 | NA | NA |
| T7RNAP_79 | NA | NA |
| T7RNAP_80 | NA | NA |
| T7RNAP_81 | NA | NA |
| T7RNAP_82 | NA | NA |
| T7RNAP_83 | NA | NA |
| T7RNAP_84 | NA | NA |
| T7RNAP_85 | NA | NA |
| T7RNAP_86 | NA | NA |
| T7RNAP_87 | NA | NA |
| T7RNAP_88 | NA | NA |
| T7RNAP_89 | NA | NA |
| T7RNAP_90 | NA | NA |
| T7RNAP_91 | NA | NA |
| T7RNAP_92 | NA | NA |
| T7RNAP_93 | NA | NA |
| T7RNAP_94 | NA | NA |
| T7RNAP_95 | NA | NA |
| T7RNAP_96 | NA | NA |
| T7RNAP_97 | NA | NA |
| T7RNAP_98 | NA | NA |
| T7RNAP_99 | NA | NA |
